# Pervasive translation of circular RNAs driven by short IRES-like elements

**DOI:** 10.1101/473207

**Authors:** Xiaojuan Fan, Yun Yang, Chuyun Chen, Zefeng Wang

## Abstract

Although some circular RNAs (circRNAs) were found to be translated through IRES-driven mechanism, the scope and functions of circRNA translation are unclear because endogenous IRESs are rare. To determine the prevalence and mechanism of circRNA translation, we developed a cell-based system to screen random sequences and identified 97 overrepresented hexamers that drive cap-independent circRNA translation. These IRES-like short elements are significantly enriched in endogenous circRNAs and sufficient to drive circRNA translation. We further identified multiple *trans*-acting factors that bind these IRES-like elements to initiate translation. Using mass-spectrometry data, hundreds of circRNA-coded peptides were identified, most of which have low abundance due to rapid degradation. As judged by mass-spectrometry, 50% of translatable endogenous circRNAs undergo rolling circle translation, several of which were experimentally validated. Consistently, mutations of the IRES-like element in one circRNA reduced its translation. Collectively, our findings suggest a pervasive translation of circRNAs, providing profound implications in translation control.

## Introduction

Circular RNAs (circRNAs) have recently been demonstrated as a class of abundant and conserved RNAs in animals and plants (for review, see ^1–3^). Most circRNAs are produced from a special type of alternative splicing known as back-splicing, and are predominantly localized in cytoplasm ^3–5^. However, the general function of circRNA *in vivo* is still an open question. Several circRNAs have been reported to function as molecular sponges to sequester miRNAs ^6, 7^ or RNA binding proteins (RBPs) ^8^ (i.e., as competitors of the linear mRNAs), whereas some nuclear circRNAs were reported to promote transcription of nearby genes ^9, 10^. Since *in vitro* synthesized circRNAs can be translated in cap-independent fashion ^11^ and most circRNAs are localized in cytoplasm, it is highly possible that many circRNAs function as mRNAs to direct protein synthesis.

Recently we and other groups reported that some circRNAs can indeed be translated *in vivo* via different internal ribosome entry sites (IRESs) ^12–15^. Because circRNAs lack a 5≠ end, the translation of circRNAs can only be initiated through a cap-independent mechanism that requires the internal ribosomal entry site (IRES). However the endogenous IRESs are infrequent in eukaryotic transcriptomes, and even their existence is sometimes under debate ^16–18^, which casted doubts on the scope of circRNA translation. In support of this notion, a recent study has identified hundreds of putative IRESs by systematically searching selected viral sequences and 5≠-UTR of human mRNA ^19^, however only a small fraction (<1.5%) of ∼100,000 known circRNAs ^20^ contain these newly identified IRESs.

To study the scope of circRNA translation, we developed a cell-based reporter system to screen a random library for short sequences that drive circRNA translation. Through a near-saturated screen and subsequent bioinformatics analyses, we identified 97 IRES-like hexamers that can be clustered into 11 groups with AU rich consensus motifs. The IRES-like activities of these short motifs were further validated experimentally. Importantly, the IRES-like elements are significantly enriched in human circRNAs compared to all linear RNAs, suggesting that they are positively selected in circRNAs. Since these IRES-like hexamers account for ∼2% of all hexamers (97/4096), any sequences longer than 50-nt should contain such an element by chance, implying that most circRNAs in human cells can potentially be translated through the IRES-like short elements. Consistently, we found that circRNAs containing only the coding sequences can indeed be translated in a rolling circle fashion, presumably from internal IRES-like short elements in the coding region. We further identified hundreds of circRNA-coded proteins with mass spectrometry datasets, and explored the potential roles of the circRNA-coded proteins and the mechanism of their translation. Collectively, our data indicate that short IRES-like elements can drive extensive circRNA translation, which may represent a general function of circRNAs.

## Result

### Unbiased identification of short IRES-like elements

Previously we reported that GFP-coding circRNAs can be translated from different viral or endogenous IRESs ^12, 13^. Similar circRNA translation reporters were also used by other researchers ^21^, and were also previously validated using an *in vitro* translation assay ^22^. Surprisingly, we found that three of the four short poly-N sequences used as negative controls for known IRESs were also found to promote GFP translation, with the only exception of poly-G (Fig. S1A). This observation indicates that certain short elements other than known IRESs are sufficient to initiate circRNA translation. To systematically identify additional sequences that drive circRNA translation, we adopted an unbiased screen approach originally developed to identify splicing regulatory *cis*-elements ^23–25^. Briefly, a library of random 10-nt sequences was inserted before the start codon of circRNA-coded GFP ^26^, which was transfected into 293T cells to generate circRNAs that can be translated into intact GFP (see Methods and Table S1). The cells with active circRNA translation (i.e. green cells) can be recovered with fluorescence activated cell sorting (FACS), and the inserted decamers can be subsequently sequenced to identify the IRES-like elements that drive circRNA translation.

To achieve full coverage of entire library, >100 million cells were transfected (Fig. 1A). We sequenced the inserted fragments from both dark and green cells using high-throughput sequencing, and compared the resulting decamers between these two cell populations to extract the hexamers enriched in the green cells *vs.* the dark cells (Fig. 1B, Fig. S1B). We have identified 97 hexamers that are significantly enriched in the cells with GFP fluorescence (Table S2). These enriched sequences are generally AU-rich despite the pre-sorting library has roughly even base composition (Fig. 1C and Fig. S1C). In addition, these hexamers have strong dinucleotide biases toward AC, AG, AT and GA (Fig. 1D). Based on sequence similarity, the 97 hexamers enriched in green cells were further clustered into 11 groups to produce consensus motifs (Fig. 1E, top). Most consensus motifs are AU-rich motifs and enriched in 3≠-UTR of linear mRNA (Fig. S1D). Consistent with the previous report that m6A modification sites can function as IRES to drive circRNA translation ^13^, several enriched hexamers contain the RRACH signature for the m6A modification, however they were not prevalent enough to be clustered into a consensus motif.

**Figure 1.**
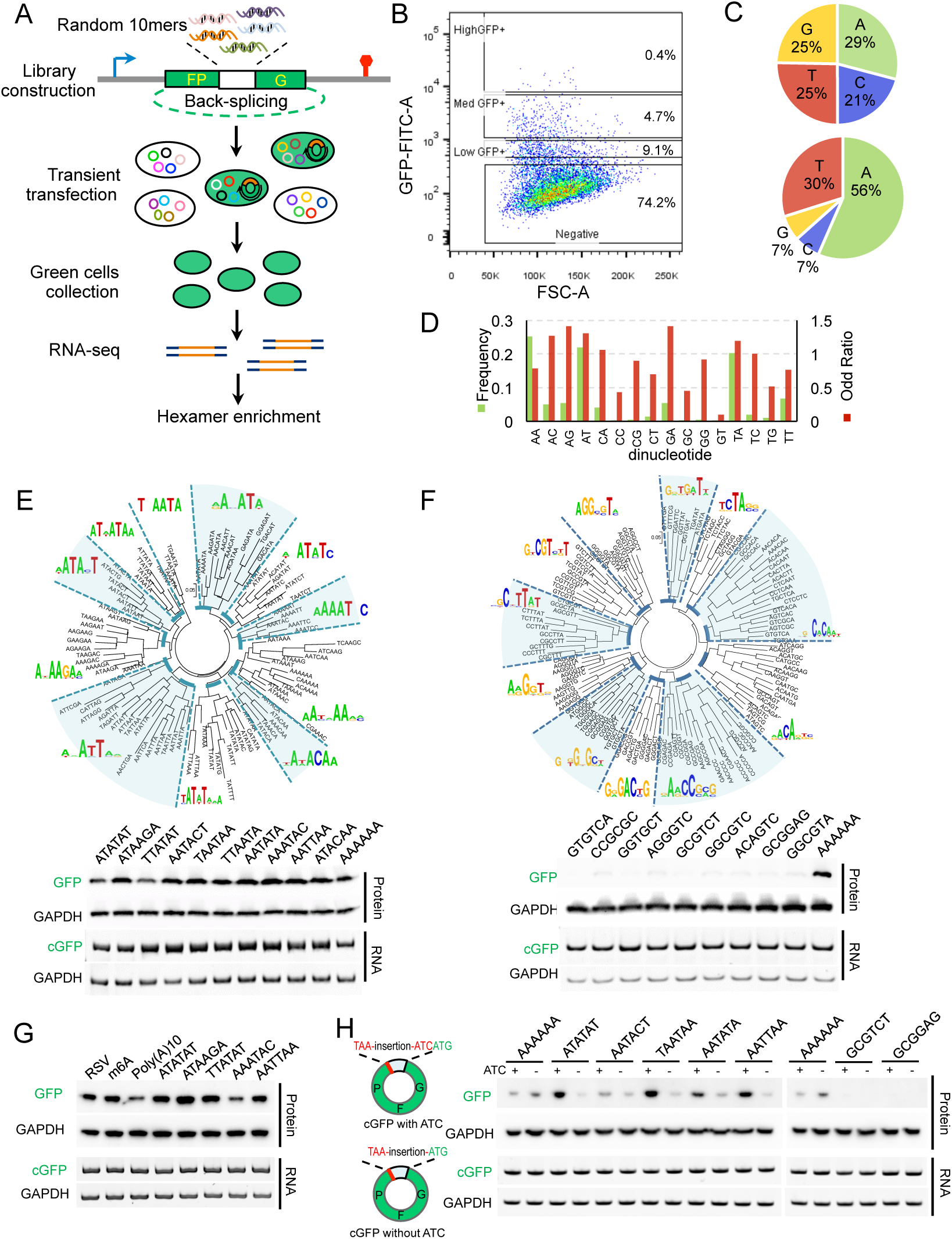
Extensive IRES-like elements can drive circRNA translation. **(A)** Schematic diagram for screening short IRES-like elements. Random decamers were inserted into pcircGFP-BsmBI reporter, and the resulting library was transfected into 293T cells and sorted by FACS. The green cells were collected and the inserted sequences were sequenced using high-throughput sequencing. The hexamers enriched in green cells were identified by computational analysis (also see method). **(B)** Flow-cytometry analysis of cells transfected with circRNA reporter containing the random 10-mer library. The cells were classified into four groups based on their GFP fluorescence (GFP negative cells and cells with low, medium or high GFP signals). The cells with medium and high fluorescence were sorted as “green cells”. **(C)** Single nucleotide frequency in the starting library (top) and the sequences enriched in green cells (bottom). **(D)** Frequencies and odd ratios of the dinucleotide in the sequences enriched in green cells. The odd ratio is defined as the probability of a dinucleotide divided by the product of the probabilities of each base of the dinucleotide. **(E)** The 97 enriched hexamers (i.e., IRES-like elements, z score >7) were clustered into 11 groups with the consensus motifs shown as pictogram (top). The representative hexamers in each cluster were inserted back into the circRNA reporter, and the resulting reporters that were transiently transfected into 293T cells. The translation of GFP was assayed by western blot at 48 hours after transfection (bottom). **(F)** The 122 depleted hexamers (i.e., negative control, z score < −7) were clustered into 11 groups and the consensus motifs were shown as pictogram (top). The activities of representative hexamers in all clusters were tested using the same condition as panel E. An IRES-like element, AAAAAA, was included as the positive control. **(G)** Comparison of newly identified IRES-like elements with m6A sites for the activity to drive circRNA translation. Two short sequences containing m6A sites (RSV and m6A) and several IRES-like elements were tested using the same condition as panel E. **(H)** Effects of Kozak sequence on circRNA translation. The enriched and depleted hexamers were inserted into circRNA reporters with or without ATC trimer (half Kozak sequence), and were transfected into 293T cells. The samples were analyzed using the same condition as panel E. The circRNA reporters inserted with poly-A sequence were loaded twice as control in both blots.

The IRES-like activities of each cluster were further validated by inserting the representative hexamers (or the control hexamers depleted in green cells) into the circRNA reporter to examine the translation product of the resulting circRNAs (Fig. 1E, bottom). All the reporters inserted with the enriched hexamers showed robust GFP translation from circRNAs, whereas the GFP productions from the reporters inserted with control hexamers were barely detectable (Fig. 1F). Furthermore, the enriched hexmers showed comparable activities as two short m6A-containing sequences that were previously reported to drive circRNA translation (Fig. 1G) ^13^. In addition, the circRNA levels were similar in all reporters containing different sequences as judged by RT-PCR and by northern blot (Fig. 1E-G and Fig. S1E), suggesting that the differences in GFP production are probably due to distinct activities of these hexamers in driving translation rather than by differences in back-splicing efficiency..

It is well accepted that “Kozak” sequence can improve the mRNA translation efficiency ^27^, and our circRNA reporters contain a Kozak-like sequence (ATC) before the start codon. To examine if the activity of the enriched hexamers is independent from the Kozak-like ATC sequence, we generated a mutated reporter by deleting the Kozak-like “ATC”. We found that all enriched hexamers tested can still direct circRNA translation from the reporters with “ATC” deletion, whereas no translation was observed from the depleted hexamers (Fig. 1H), suggesting that these enriched hexamers can drive circRNA translation in the absence of Kozak-like sequence. Interestingly, the deletion of “ATC” sequence reduced translation in 4 out of 6 enriched hexamers tested but increased translation driven by one hexamer (poly-A sequences). These results suggested that circRNA translation by enriched hexamers can be improved by the Kozak sequence in some cases, but the Kozak sequence is not absolutely required.

We further examined the IRES-like activity of the enriched hexamers using *in vitro* synthesized circRNA by self-splicing of group I intron ^28^, which were further purified using HPLC for transfection (Fig. S1F). Compared to the controls, all tested circRNAs with candidate IRESs can be translated into luciferase in two cell lines (Fig. S1G), and the newly identified short IRES-like elements have similar activity as known IRESs of endogenous genes (5’UTR from Hsp70, CCGGCGG from Gtx, and UACUCCC from OR4F17) ^29, 30^. However, the viral IRESs from EMCV and CSFV have a higher activity in such *in vitro* system, probably because their activities are less dependent on the nuclear RNA binding proteins or modifications of the *in vitr*o synthesized RNAs ^31, 32^. Collectively, these results indicated that our screen can reliably identify short sequences that drive circRNA translation, and thus we refer this set of 97 hexamers as IRES-like hexamers.

### IRES-like hexamers are enriched in endogenous circRNAs to drive translation

We further examined the frequency of each hexamer in linear mRNAs and circRNAs, and compared the average frequency of the IRES-like hexamers *vs.* the control set of hexamers in different types of RNAs. In the linear mRNAs (all annotated mRNAs from RefSeq), the average frequencies of the IRES-like hexamers were similar to the random hexamers or the hexamers depleted in green cells (Fig. 2A, left panel). Surprisingly, the average frequencies of IRES-like hexamers were significantly higher in all tested circRNA datasets ^4, 7, 33, 34^ compared to the control hexamer sets (Fig. 2A), indicating that endogenous circRNAs are enriched with short IRES-like elements. Since these IRES-like hexamers were independently identified from unbiased screen of random sequences, such enrichment strongly suggests that circRNAs may be positively selected for their ability to be translated.

**Figure 2.**
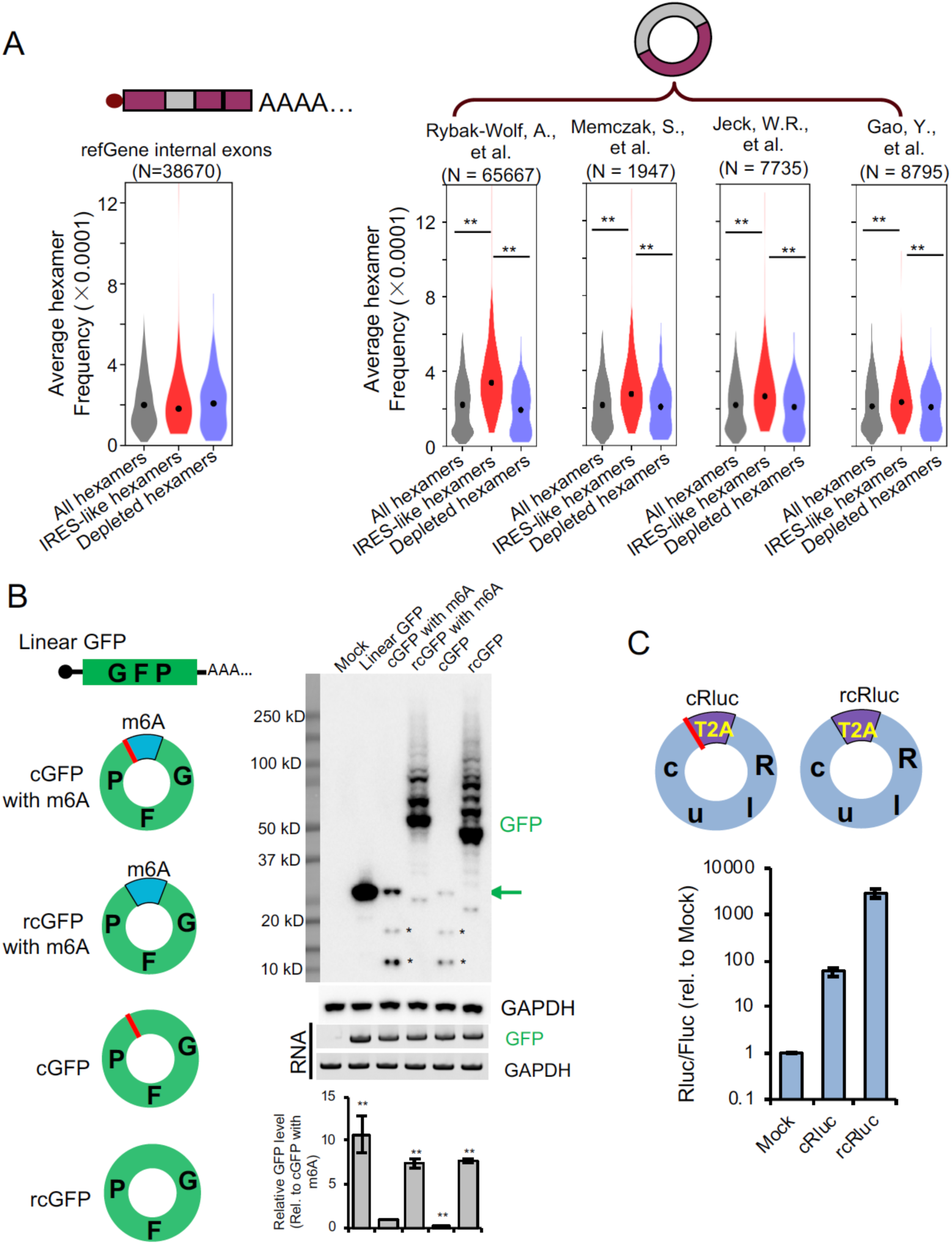
circRNAs contain many IRES-like elements that initiate translation. **(A)** IRES-like elements are significantly enriched in circRNAs. Average frequencies of different types of hexamers (all hexamers, IRES-like hexamers and depleted hexamers) in the internal exons of linear mRNAs *vs.* circRNAs were plotted. N: the number of RNA sequences in each dataset. **: p-value < 0.001 with Kolmogorov– Smirnov test. **(B)** Translation of circRNAs can be initiated by internal coding sequence. Left panel, schematic diagrams of expression reporters for a linear mRNA and four circRNAs that code for GFP. From top to bottom: linear GFP mRNA; circRNA with an m6A site at upstream of the start codon; circRNA with an upstream m6A site and no stop codon; circRNA with start codon immediately following the stop codon; circRNA containing only the coding sequence (no stop codon or UTR). Red line, stop codon. The circRNA plasmids were transfected into 293T cells, and samples were analyzed by western blot at 2 days after transfection (right panel). The level of GAPDH was measured as a loading control. The green arrow indicates the full-length GFP, and the asterisks indicate the truncated GFP proteins translated using internal start codons. The bar graph represents the quantification of GFP protein levels relative to GAPDH. The protein levels were also normalized to the RNA (n = 3, mean ± SD, **: p-value < 0.01 with Student’s t test comparing to the cGFP with m6A). **(C)** Translation of the circRNA-coded Renilla luciferase (Rluc) using internal IRES-like elements. Top: schematic diagram of two Rluc circRNAs, where the Rluc ORF was split into two parts so that the full-length Rluc ORF can only be generated through back-splicing. The cRluc contains a sequence coding a T2A peptide by which the translation product can be cleaved into full length Rluc protein. rcRluc does not contain stop codon. Bottom: dual luciferase assay of cRluc and rcRluc. Control and circular Rluc plasmids were co-transfected with Fluc (firefly luciferase) reference reporter into 293T cells. The cells were lysed at 48 hours after transfection for luminescence measurement using luminescence reader, and the relative luminescence signals were plotted (mean ± SD, n = 3 independent experiments).

The 97 IRES-like hexamers account for ∼2% of the entire hexamer population (4^6^=4096), indicating that there will be an IRES-like hexamer in a 50-nt sequence by chance. Since >99% circRNAs are longer than 100-nt ^20^ (Fig. S2A), most circRNAs should contain internal IRES-like short elements by chance. Therefore, almost all open reading frames (ORFs) in circRNAs could potentially be translated using such IRES-like short elements. To directly test this surprising conclusion, we generated a series of GFP-coding circRNA reporters without known IRES sequences or stop codon to measure GFP production using western blot (Fig. 2B). Consistent with our previous report, the control reporter containing the m6A modification sites at upstream of the start codon was reliably translated ^13^, and deletion of stop codon in this reporter led to production of GFP concatemers through the rolling circle translation of circRNA (Fig. 2B). When deleting all untranslated sequences between the start and stop codon in the circRNA, the intact GFP translation is abolished, presumably because there is no room for any sequences to function as the IRES. However, when deleting the stop codon, the circRNA containing only the GFP coding sequence can also be translated in a rolling circle fashion to produce GFP concatemers, presumably through an internal sequence function as an IRES (lanes 4 and 5, Fig. 2B). As a control, we also confirmed the efficient circRNA expression in all the RNA samples using northern blot with the optional RNase R treatment (Fig. S2B).

Interestingly, the rolling circle translation can produce some huge GFP concatemers (lanes 3 and 5, Fig. 2B, the huge proteins could not be efficiently transferred to the membrane and thus being underestimated). To eliminate the possible artifacts from rolling circle transcription of circular plasmid, these reporter plasmids were stably inserted into genome using Flp-In system, and the similar rolling circle translations were observed in these stably transfected cells (Fig. S2C).

In addition, the rolling circle translation initiated from internal coding sequence is not limited to the GFP gene, as the circRNAs containing only the ORF sequences of different luciferase genes can also be translated from internal coding sequence (Fig. 2C, Fig. S2D-F). Interestingly, the rolling circle translation from circRNAs apparently produced more proteins than the circRNAs with stop codons (Fig. 2B), suggesting that initiation of circRNA translation may be the rate-limiting step as the ribosome recycling and reinitiation is unnecessary for rolling circle translation ^22, 35, 36^.

### *Trans*-acting factors that bind to IRES-like short elements to promote circRNA translation

With the identification of multiple IRES-like short elements, we next seek to determine the molecular mechanisms by which these elements initiate cap-independent translation of circRNA. An earlier report showed that short sequences may function as IRES by pairing with certain regions of 18S rRNA (i.e. active region) ^19^. However, we found little correlation between our newly identified IRES-like elements to these “active 18S rRNA regions” (Fig. S3A), suggesting that these newly identified elements may not function by paring with 18S rRNA.

Previously we found that m6A reader protein YTHDF3 can recognize N6-methyladenosine in circRNA to directly recruit translation initiation factors ^13^. By analogy we hypothesized that the newly identified IRES-like short elements may also function as regulatory *cis*-elements to recruit *trans*-acting factors that promote translation ^37^. To identify such *trans*-acting factors, we used the consensus sequences of IRES-like elements as bait for affinity purification of their specific binding proteins (Fig. 3A). Briefly, the chemically synthesized 20-nt single-strand RNA oligonucleotides containing three copies of IRES-like hexamers and a 5≠ end biotin modification were incubated with HeLa cell extracts, and the RNA-protein complexes were purified with streptavidin beads ^23–25^ (Fig. 3A). We found that, all the five RNA probes consist of IRES-like elements showed robust binding of several proteins, whereas the negative sequence (ACCGCG) had weak background of non-specific RBP binding (Fig. 3B). The specific protein bands in each lane were collected and subsequently analyzed with mass-spectrometry (LC-MS/MS), and the top candidates in each band were identified as candidate *trans*-acting factors (Fig. 3B).

**Figure 3.**
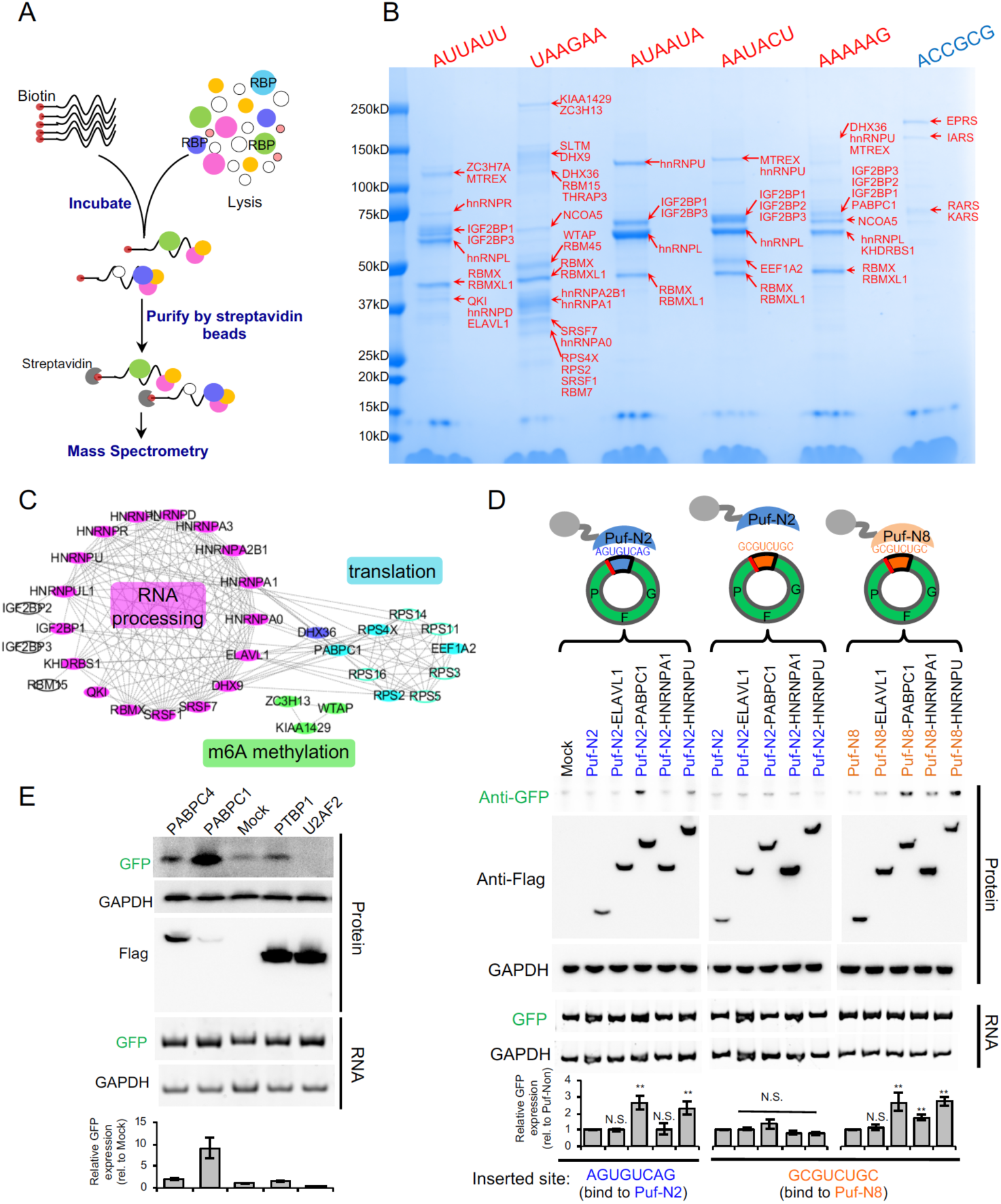
Systematically identification of *trans*-factors that recognize IRES-like elements. **(A)** Schematic diagram of RNA affinity purification. Biotin-labeled RNAs containing consensus motifs of IRES-like elements were incubated with HeLa cell lysate, and RNA-protein complexes were purified by streptavidin beads. The proteins were further identified by mass spectrometry (see method for details). **(B)** Identification of *trans*-factors bound by each RNA probe. The probes presenting five consensus motifs of IRES-like elements (red) and a control probe (blue) were used (see table S2 for full sequence). The total proteins eluded from each RNA probe were separated with SDS-PAGE, and each band was cut and analyzed by mass spectrometry. The top three identified proteins in each band were labeled at right in red. **(C)** Protein-protein interaction network of identified *trans*-factors. Top proteins bound by all RNA probes (i.e., IRES-like elements) were analyzed by STRING and clustered into two main groups by MCODE tool. **(D)** Measurement of the activity of *trans*-factors. The circRNA reporter inserted with two depleted hexamers in tandem (with very weak IRES-activity by itself) were co-transfected into 293T cells with different Puf-fusion proteins that specifically recognize an 8-nt target in the inserted sequences. The resulting cells were collected at 2 days after transfection to analyze the protein and RNA levels by western blot and RT-PCR respectively. Different pairs of Puf proteins and 8-nt targets were used as specificity control. Puf-N2 can specifically bind AGUGUCAG, whereas Puf-N8 can specifically bind to GCGUCUGC. The bar graph represents the quantification of GFP levels relative to GAPDH. The protein levels were also normalized to the RNA (n = 3, mean ± SD, **: p-value < 0.01 with Student’s t test comparing to the control sample, N.S.: not significant). **(E)** Validation of PABPC1 activity. The expression vector of PABPC1 and various control RBPs were co-transfected with circRNA translation reporter containing (A)_10_ sequence before the start codon, and the protein products were assayed at 48 hours after transfection. The bar graph represents the quantification of GFP levels relative to GAPDH. The protein levels were also normalized to the RNA (n = 3, mean ± SD).

In total 58 protein candidates were identified with high confidence, with many overlapping proteins in different baits (Table S3). The majority of these proteins are known to bind RNAs and can be clustered into two major groups based on the protein-protein interaction (PPI) network: the proteins involved in RNA processing (e.g., hnRNPs) and the proteins involved in mRNA translation (e.g., the ribosomal proteins) (Fig. 3C). The identified RBPs are enriched for regulatory function in RNA processing, splicing, transport and stabilization as judged by gene ontology analysis (Fig. S3B).

To validate the function of these proteins, we fused the candidate RBPs to a programmable RNA binding domain (i.e., Puf domain) that can be designed to bind any 8-nt RNA sequences ^38^, and co-expressed the fusion proteins with the circRNAs containing their cognate targets. We found that the specific tethering of PABPC1 and hnRNP U clearly promoted translation of the circRNA, whereas the ELAVL1 (HuR) and hnRNP A1 did not affect translation when tethered to the same site (Fig. 3D). Such translation promoting activity required the specific binding of *trans*-acting factors to circRNAs, as disrupting the Puf-RNA interactions had abolished the regulatory effect and restoring the specific interaction can rescue the translation-promoting activity (Fig. 3D).

PABPC1 is an abundant protein that binds to poly-A or AU-rich sequences ^39^. We further examined the role of PABPC1 in regulating circRNA using reporters contain short poly-A elements at upstream of start codon in circRNA (Fig. 3E). The results showed that over-expression of PABPC1 can indeed promote translation of circRNA-coded GFP. As controls, the circRNA translation was not affected by another poly-A binding protein PABPC4 or by PTBP1 that was previously reported to enhance IRES activity (Fig. 3E). Interestingly, for unknown reason, the co-expression of U2AF2 seemed to inhibit translation of the circRNA (Fig. 3E). Taking together, our results showed that certain RBPs (e.g., PABPC1) are capable to recognize these IRES-like elements to promote cap-independent translation of circRNAs, exemplifying a new model for circRNA translation initiation independent of canonical IRESs.

### Identification of circRNA-coded proteins

Since most circRNAs can potentially be translated, we further examined molecular characteristic of the putative circRNA-coded proteins. In accordance with previous observation ^15^, we found that the exons in the 5≠ end of a pre-mRNA are more likely to be included in the circRNAs (Fig. 4A), suggesting that many circRNAs could potentially code for N-terminal truncated protein isoforms of their host genes. Consistently we found that 79% of circRNA exons are from coding region, while 17% of circRNA exons spanning 5≠-UTR and coding region and 4% spanning coding and 3≠-UTR region (Fig. 4A inset).

**Figure 4.**
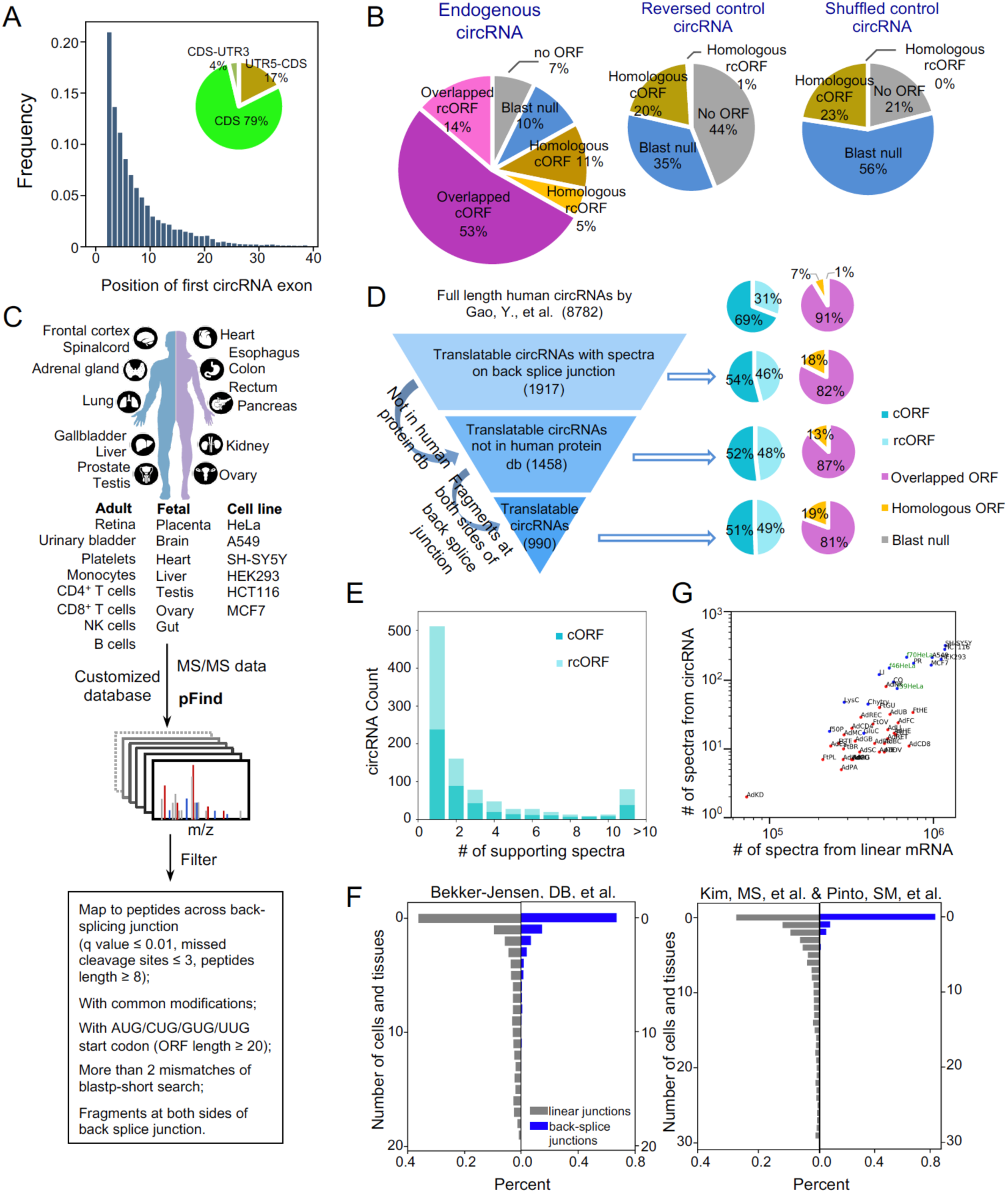
**Identification of circRNA-coded proteins.** (A) Position distribution for the first exon of circRNAs in their host genes. Full length circRNA sequences were analyzed based on previously published datasets, and the histogram was plotted according to the position of the first circRNA exon (i.e., the exon number) in host genes. The inserted pie chart presents the percent of circRNAs overlapping with different regions of mRNA. (B) Survey of the potential coding products of circRNAs. Left panel, percent of endogenous circRNAs that code for an ORF longer than 20 aa. Purple pie slices: circRNAs translated in a regular fashion (cORF, dark purple) or a rolling circle fashion (rcORF, light purple) into proteins that are partially overlapped with their host genes; brown pie slices: circRNAs translated in a regular fashion (cORF, dark brown) or a rolling circle fashion (rcORF, light brown) into proteins that are not overlapped with their host genes but are homologous to other known proteins; blue pie slices: circRNAs translated into proteins that are not homologous to any known proteins (cORF and rcORF combined); grey pie slices: circRNAs do not contain any potential ORF longer than 20 aa. Right panels: the same analysis of putative coding products from two control circRNA sets, reversed sequences of endogenous circRNAs and random shuffled sequences of endogenous circRNAs. (C) Schematic diagram for identification of circRNA-coded proteins using proteomic datasets. (D) Schematic diagram of translatable circRNA identification pipeline. Left, computational filters sequentially applied to identify translatable circRNAs, the numbers of circRNAs passing each filter. Right, the percentage of different types of circRNA-coded ORFs (rcORFs, cORFs, overlapped ORFs, homologous ORFs, and blast null) in total circRNAs that passed each filter. The definition of different types of ORFs encoded by circRNAs are same as panel B. (E) Distribution of the supporting spectra for each translatable circRNA. (F) Distribution of the number of cell lines and tissues for each translatable circRNA in two proteomic datasets. (G) Comparison of the numbers of spectra from linear mRNAs vs. circRNAs. Abbreviation of different tissues and cell lines are listed in table S4. Blue dot: proteomic data from Bekker-Jensen, DB, et al; Red dot: proteomic data from Kim, MS, et al. & Pinto, SM, et al.. Green words indicate the 39 fractions, 46 fractions and 70 fractions from HeLa cells using high-capacity offline HpH reversed-phase LC.

Using the dataset of full length human circRNAs ^34^, we examined the putative circRNA-coded ORFs that are longer than 20 amino acids (equivalent to 60-nt). We found that a large fraction of endogenous circRNAs (67%) can code for proteins overlapping with their host genes (i.e., translated in the same reading frame as the host gene), including 14% of endogenous circRNAs that can be translated in a rolling circle fashion (named as overlapped cORF and rcORF respectively, purple pie slices in the left panel of Fig. 4B). In addition, 16% of human circRNAs can code for proteins that are different from host genes but are homologous to other known proteins (homologous cORF and rcORF, brown pie slices in Fig. 4B, left panel), whereas 10% circRNAs code for proteins that are not homologous to any known proteins on earth. In fact, only 7% of circRNAs have no ORF longer than 20 aa. In comparison, a much larger fraction of circRNAs from two different controls (reversed or shuffled sequence controls) do not contain ORFs longer than 20 aa (Fig. 4B, right two panels), and the remaining control circRNAs are more likely to code for proteins that are not homologous to any known proteins, suggesting that the endogenous circRNAs are more likely to code for functional proteins compared to controls.

To systematically identify circRNA-coded proteins, we searched raw mass spectra from publically available tandem mass spectrometry (MS-MS) datasets for possible peptides across the back splice junctions of all published circRNAs ^20, 34^ (Fig. 4C). Two sets of high-resolution comprehensive human proteomic data were selected, including raw mass spectra data from 30 tissues and 6 cell lines ^40, 41^. We used open pFind(v3.1.3) ^42^ to search against a combined database of UniProt human proteins and potential peptides encoded by back splice junctions of circRNAs. We applied fairly stringent thresholds (see Methods) to identify the peptides that are encoded only by back splice junctions of circRNAs but not found in any known proteins. Two additional filters were used to reduce false positives in our search: We discarded peptides similar to canonical proteins in non-redundant human protein database (0-2 mismatches), and required the positive spectra to contain peptide fragments encoded by both sides of back splice junction.

Our search of endogenous circRNA-coded proteins identified 2721 mass spectra across 990 back splice junctions from 646 human genes, all of which contain putative cORF or rcORF longer than 20 amino acids (Fig. 4D, Table S4). Interestingly, we found that the fraction of circRNAs with rolling circle translation products increased as the additional filters were applied in our search (∼50% of the circRNAs contain putative rcORFs at the end, see Fig. 4D). Such increase is consistent with a higher efficiency of rolling circle translation where the re-initiation is not required (Fig. 2B). Alternatively, the higher detection rate for translation products of rcORFs may partially due to an inherent bias of detecting such proteins, which contain multiple copies of the same peptide coded by the back splice junction.

More than 80% of the identified circRNA-coded peptides overlapped with the translation products of their host genes (Fig. 4D), suggesting that the circRNAs preferably produce different translation isoforms of the host genes. Interestingly, gene ontology analyses indicated that their host genes are significantly enriched with the functions in RNA translation, RNA splicing/processing, and platelet degranulation (Fig. S4), implying that many circRNAs may code for new protein isoforms with potential roles in regulating these biological processes. The enrichment in platelet degranulation may also provide a functional implication to the previous observation that the circRNAs are highly expressed in platelet ^43, 44^.

For many circRNAs, we identified multiple mass spectra to support the same back splice junctions, including 80 circRNAs with >10 different spectra across their back splice junctions (Fig. 4E). Further analyses showed that the circRNA-coded peptides are mostly presented in a small set of cells or tissues, with 60-80% of circRNA-coded peptides being identified only from a single sample in both MS-MS datasets (Fig. 4F, blue bars). In comparison, the peptides encoded by the adjacent splicing junctions from the linear mRNAs are more ubiquitously expressed, with some peptides being found in all samples (Fig. 4F, grey bars). These results suggest that the circRNA-coded proteins are generally more specific to certain cell types or tissues.

### circRNA-coded proteins have low abundance partially due to rapid degradation

We next analyzed the numbers of mass spectra supporting circRNA-coded peptides (i.e. peptides across back splice junctions) in all samples and compared to those from known proteins encoded by the canonical linear mRNAs. As expected, the numbers of newly identified circRNA-coded peptides were positively correlated with the numbers of total peptides in two independent datasets (Fig. 4G). In addition, applying additional fractionations in the same sample (e.g., 39, 46 and 70 fractions of HeLa cells) increased the numbers of newly identified circRNA-coded peptides (Fig. 4G). These results suggested that the proteomic identifications of circRNA-coded proteins have not reached saturation, and thus the spectrum numbers of supporting peptides will be roughly correlated to the abundance of these “new proteins”.

To estimate the relative abundance of circRNA-coded proteins, we compared the peptides across back splice junction *vs.* those across adjacent splicing junctions of the host linear mRNA. We found that the circRNA-coded peptides generally have a smaller number of spectra to support each peptide as compared to those from the peptides encoded by linear adjacent splice junctions (Fig. S5A), suggesting that the circRNA-coded peptides have much lower abundance than their linear counterparts. Consistently, the q values of the mass spectra supporting circRNA-coded peptides were much higher than those supporting the peptides across linear adjacent splice junctions (Fig. S5A). Since the more abundant proteins are usually supported by MS data with higher confidence, this result again suggested that the circRNA-coded proteins generally have lower abundance than their linear counterparts.

Even for the canonical proteins, we can only detect a small number of peptides across the splice junctions using MS-MS data (4254 out of 67912 adjacent canonical junctions) (Fig. S5A). By analyzing the sequence composition across splicing junctions, we found that lysine and arginine are enriched at the −1 position of all splicing junctions (Fig. S5B). Specifically, >70% of splice junctions contain at least one lysine or arginine regardless of linear or back splice junctions (data not show). Since the current proteomic samples are mostly lysed by trypsin with a cleavage site of lysine or arginine, such sequence bias leads to a significant depletion of peptides across splice junctions, which is consistent with an earlier report ^45^. Unlike the proteins encoded by linear mRNAs, the circRNA-coded proteins can only be recognized by peptides across back splice junctions, which could partially explain why the circRNA-coded peptides are difficult to identify. This result further suggested that additional circRNA-coded proteins could be identified by optimizing MS methods with different protease cleavage.

In addition to the above technical difficulties to detect circRNA-coded proteins, multiple factors may also contribute to the low abundance of circRNA-encoded proteins. First, the abundance of most circRNAs are generally known to be lower than their linear counterparts ^46–48^, which should partially contribute to the low abundance of the circRNA-coded peptides. In addition, the low abundance of circRNA-coded proteins may also due to a slow protein synthesis and/or a fast protein degradation. It is previously known that the efficiency of cap-independent translation is relatively low (Fig. 2B, also see reviews ^49, 50^). However, the stability of the circRNA-coded proteins is still an open question: Although the majority of circRNA-coded protein sequences overlap with the proteins encoded by endogenous host gene, it is possible that the extra peptides specifically encoded by sequences across the back-splicing junction make the protein unstable.

To directly address this possibility, we selected three C-terminal peptides specifically encoded by the sequences across the back splice sites of different circRNAs, and fused them to the C-terminus of GFP in the circGFP reporters (Fig. S5C). Tethering the potential degron onto a test protein (like GFP) is a routine assay to measure protein degradation ^51, 52^, with the usual caveats in selecting specific test protein. The resulting circRNA will code for a GFP fusion protein containing different circRNA-specific peptides, and we found that these GFPs with circRNA-coded tails are expressed in much lower level compared to the one without these tails (Fig. S5D). Moreover, the expression level of the fusion proteins with circRNA-coded tails increased upon the inhibition of proteasome degradation by MG132 treatment, whereas the GFP without such tails was essentially unaffected (Fig. S5E). In contrast, the levels of all GFP fusion proteins were not affected by chloroquine treatment that inhibits autophagic protein degradation (Fig. S5E). These observations indicate that the circRNA-coded C-terminal tails can indeed destabilize many circRNA-coded proteins, and such degradation was mainly mediated by proteasomes.

### Translation products derived from circRNAs

Although the endogenous circRNAs were found to contain more cORFs (with the cORF:rcORF ratio roughly equaling to 3.4, Fig. 4B), further analyses of proteomic data showed that ∼50% of the circRNA-coded peptides are derived from the endogenous circRNAs containing rcORFs (Fig. 4D), suggesting that rcORFs are more efficiently translated from circRNAs (consistent with observations in Fig. 2B and 2C). Since the endogenous circRNAs had not been reported to undergo rolling circle translation, we seek to further determine whether these identified circRNAs with rcORFs are indeed translated in different cells.

Detection of translation products of endogenous circRNAs is technically difficult because they differ from the canonical products only at the back splice junctions. Generating new antibodies against the short fragment coded by the back splice junctions is very time consuming and sometimes unreliable, and thus we labelled the candidate circRNA products with an epitope tag. We selected three circRNAs (circPSAP, circPFAS, and circABHD12) with high quality mass spectra across back splice junctions (Fig. 5A), and constructed back-splicing reporters to ectopically express these circRNAs in two different cell types ^26^. A V5 epitope tag was inserted into the same reading frame of rcORFs to facilitate the detection of the translation products by western blot (Fig. 5B). We found multiple protein bands in 293T and SH-SY5Y cells transfected with all circRNAs tested, suggesting that these circRNAs undergo robust rolling circle translation to produce protein concatemers (Fig. 5C and Fig. S6A). Interestingly, the cells transfected with circPSAP reporter produced a faint band of protein concatemer and a much stronger band corresponding to the product of a single cycle of circRNA translation, suggesting a low translation processivity for this circRNA.

**Figure 5.**
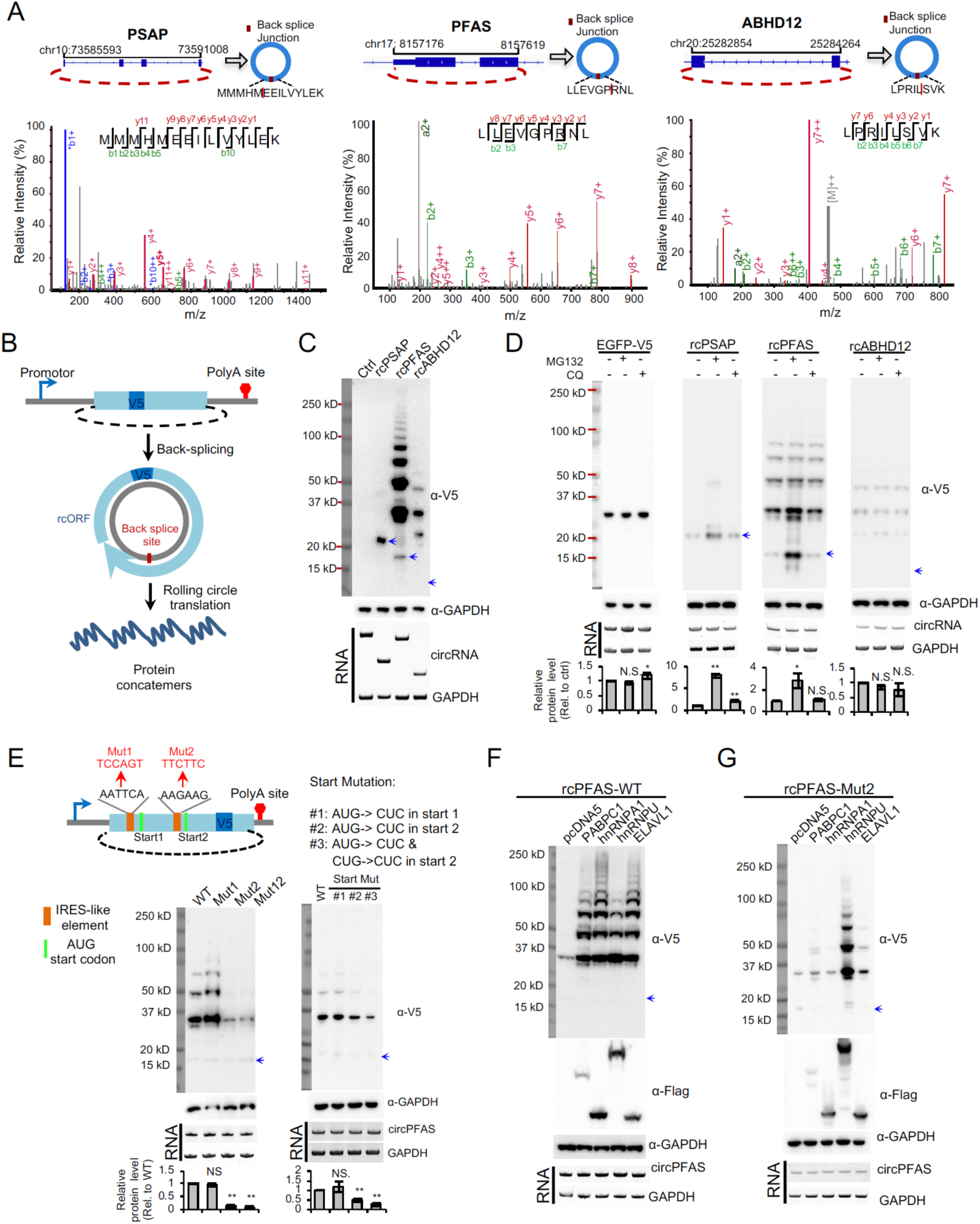
**Rolling circle translation of endogenous circRNAs.** (A) The higher-energy collisional dissociation (HCD) MS/MS spectrum of the peptide across the back splice junction of the human circPSAP (MMMHMEEILVYLEK), circPFAS (LLEVGPRNL), and circABHD12 (LPRILSVK). The annotated b- and y-ions are marked in red and green color, respectively. (B) Schematic diagram of rcORF translation reporters. The coding region of the endogenous rcORF was inserted into a back-splicing reporter. To detect the translation products from endogenous rcORF, a V5 epitope tag was inserted into the same reading frame of the rcORF. After back-splicing, the circRNA containing a rcORF and an in-frame V5 epitope tag was translated through rolling circle fashion to produce the protein concatemers. (C) The back-splicing reporters containing three endogenous rcORFs were transfected into 293 cells, the cells were collected at 48 hours after transfection, and the levels of circRNA-coded proteins and circRNAs were detected by western blotting and RT-PCR, respectively. The blue arrows represent the predicted molecular weight of the single cycle of translation product derived from cPSAP (22.4kD), cPFAS (15.6kD), or cABHD12 (10.5kD). (D) The translation products of rolling circle translation were degraded through proteasome pathway. The rcORF translation reporters were transfected into 293T cells. The transfected cells were treated with 10 µM MG132 for 2 hours, or 10 µM chloroquine for 4 hours before cell collection. The bar graph represents the quantification of protein levels relative to GAPDH. The protein levels were also normalized to the RNA (n = 3, mean ± SD, **: p-value < 0.01 with Student’s t test comparing to the untreated sample, N.S.: not significant). (E) The translation of rcPFAS was reduced by mutations on IRES-like elements or the start codon of the circPFAS. The circPFAS contains two IRES-like hexamers (AATTCA and AAGAAG), which were mutated into neutral sequences (mut1 and mut2). The downstream AUG start codons were mutated into CUC (Start Mut #1 and #2), and the non-canonical start codons CUG at downstream of the AAGAAG hexamers were further mutated into CUC (Start Mut #3). The effects of these mutations on protein production were determined with western blotting using similar procedure as described in panel C. The relative change of proteins was quantified and represented with bar graphs (n = 3, mean ± SD, **: p-value < 0.01 with Student’s t test comparing to the wild type, N.S.: not significant). (F) Co-expression of *trans*-acting factors increased the rolling circle translation products. The back-splicing reporter of rcPFAS was co-transfected into 293T cells with the expression vectors of various *trans*-acting factors that bind to the newly-identified IRES-like elements. The cells were collected and analyzed using same procedures as described in panel C. (G) The circRNA with mutated IRES-like hexamer AAGAAG (Mut2) was co-expressed with same set of *trans*-acting factors, and the production of rolling circle translation were measured using same experimental conditions described in panel F.

Since the translation of rcORFs can generate protein concatemers with repeat sequences, they are likely to induce mis-folding and aggregation of proteins, which often leads to rapid protein degradation. To measure the stability of these proteins translated from rcORF, we transfected cells with these circRNA reporters and treated the cells with short exposure of MG132 or chloroquine. We found that the translation products of two circRNAs (circPSAP and circPFAS) were increased by MG132 treatment but not by chloroquine in two different cells (Fig. 5D), whereas the ectopically expressed control protein GFP was not affected, suggesting that some rolling circle translation products were rapidly degraded in cells through proteasome pathway. These findings are consistent with the observation that the circRNA-coded proteins are generally low abundant inside cells (Fig. S5). Intriguingly, the translation product of circABHD12 seems to be relatively stable despite containing repetitive sequences (i.e., not affected by brief treatment with MG132 or chloroquine).

Next, we seek to examine whether the rolling circle translation of endogenous circRNAs is indeed driven by the newly identified IRES-like short elements. We selected the circPFAS that contains two IRES-like hexamers, and made individual mutations on both hexamers to examine if such mutations can affect its translation (Fig. 5E left, the other two circRNAs contain 3, and 10 IRES-like hexamers and thus were not tested). We found that the mutation on one of the IRES-like hexamers (AAGAAG) dramatically reduced the translation of circPFAS, whereas the mutation on the other element (AATTCA) had no effect on translation, suggesting that the AAGAAG hexamer is the major translation initiation element in circPFAS (Fig. 5E left and Fig. S6B). Interestingly, residue amount of translation product was observed even with mutations on both IRES-like elements, suggesting that other unknown *cis*-elements may also help to promote circPFAS translation. This observation is not a surprise since we used a strong cutoff (enrichment z score >7) in the identification of the 97 IRES-like hexamers, and thus may miss some elements with weak IRES-like activity. We further mutated the downstream AUG start codons to confirm that the second start codon at downstream of AAGAAG hexamer was the major translation initiation of circPFAS, as the mutation significantly reduced the translation efficiency (Fig. 5E right). Interestingly, the circRNA translation is reduced but not completely abolished with the AUG mutation, and additional mutations of nearby in-frame CUG site can further reduce the translation products (Fig. 5E). This result suggests that the cap-independent translation driven from IRES-like hexamers can also use non-canonical start codon (e.g., CUG), which is consistent with previous reports ^53^.

We next examined if the rolling circle translation of endogenous circPFAS can also be affected by the *trans*-acting factors that promote cap-independent translation of circRNA reporters (Fig. 3D and 3E). To this end, we co-expressed the circPFAS with PABPC1, hnRNP A1, hnRNP U, or ELAVL1 in two different cell types (293T and SH-SY5Y cells), and measured the products of rolling circle translation with western blot. We found that all tested *trans*-acting factors significantly increased the rolling circle translation products from circPFAS (Fig. 5F and Fig. S6C). This observation suggest that the same set of *trans*-acting factors identified earlier (Fig. 3) may also promote the rolling circle translation of endogenous circRNA, and that the translation products of a single circRNA may be affected by multiple *trans*-factors. When co-expressing these *trans*-factors with the circPFAS containing mutation on its IRES-like element (AAGAAG), we found that the translation enhancing activities of PABPC1, hnRNP A1 and ELAVL1 were dependent on the IRES-like hexamer in circPFAS, whereas the effect of hnRNP U is independent on this IRES-like element (Fig. 5G and Fig. S6D). This deference is probably because these factors bind to the IRES-like hexamer with distinct specificities and affinities, or due to subtle differences of the mechanism by which various factors affect circRNA translation.

## Discussion

Several recent studies indicated that some circRNAs can function as template to direct protein synthesis, however the nature of circRNA translation is still under debate because other studies failed to detect the significant association of circRNA with polysomes ^4, 54, 55^. In addition, the mechanism of circRNA translation is not clear. While the cap-independent translation of circRNA generally requires a viral or endogenous IRES, we have demonstrated that a short element containing m6A modification site is sufficient to drive circRNA translation by directly recruiting initiation factors ^13^. Here we surprisingly found that the requirement of IRESs can be easily fulfilled, and many short sequences (∼2% of all hexamers) are capable of driving cap-independent translation in circRNAs. In fact, any circRNAs longer than 50-nt should contain an IRES-like hexamer by chance, and more importantly endogenous circRNAs are enriched with such IRES-like elements compared to linear mRNAs. This finding is also consistent with an earlier study of *in vitro* circRNA translation, where circRNAs with mutated Kozak sequences can still be translated ^22^. As a result, thousands of cytoplasmic circRNAs can be potentially translated, among which hundreds of circRNA-coded peptides were supported by mass spectrometry evidences. We further identified many RNA binding proteins that can specifically recognize these IRES-like short elements and function as *trans*-factor to promote cap-independent translation, providing a new paradigm on the mechanism of circRNA translation.

The high sensitivity of circRNA translation reporter makes it possible to screen for short IRES-like elements. We also examined the IRES activity using a traditional bicistronic linear reporter with two tandem ORFs encoding firefly and Renilla luciferases ^56^. We found that both the known short IRESs from endogenous genes (CCGGCGG from Gtx gene, and UACUCCC from OR4F17) and the short IRES-like elements tested have little, if any, activity in direct cap-independent translation in a standard bicistronic vector (Fig. S7A), although they all worked well in circRNA reporters (Fig. S7B). The only exception is the long IRES sequence from Hsp70 that functioned in both reporters. A possible reason for the low sensitivity of the bicistronic reporter is that the linear mRNAs have higher translation background for empty control than circRNAs (∼400 fold higher, Fig. S7C), reducing its sensitivity of detection. Consistently, several problems of canonical bicistronic vectors were previously reported to produce high background due to various artifacts, including cryptic promoters, cryptic splicing sites, ribosome shunting, or re-initiation ^16–18, 57^. As a result, many data using such system have been challenged ^17, 18^. Since the circRNAs have no free 5’ or 3’ ends and can only be translated from IRES, the circRNA system may be a more reliable system for future test of IRES activity.

The majority of the identified translatable circRNAs code for new protein isoforms overlapping with canonical gene products, 50% of which can produce protein concatemers through a rolling circle translation. This finding raises an interesting open question regarding the biological function of the circRNA-coded protein isoforms. Because these circRNA-coded isoforms have large overlap with the canonical host genes, the circRNA-coded isoforms may function as competitive regulator of canonical isoforms or play the similar function in different subcellular location. On the other hand, the protein concatemers translated from circRNAs may also function as scaffold for assembly of large complexes, or form protein aggregations that are toxic to cells.

Compared to linear mRNAs, a relatively small number of circRNAs are reported to be associated with polysomes ^54, 55^, suggesting an inefficient translation of circRNAs. In addition, the minimal requirement for the cap-independent translation of circRNAs and the difficulties to detect circRNA-coded products presented an intriguing paradox. While there is no clear explanation, several factors may help to reconcile such contradiction. First, although most circRNAs have the capacity to be translated, it does not necessarily mean that they are indeed translated efficiently *in vivo* to produce stable proteins. It is possible that some products from pervasive translation of circRNAs may not be folded correctly and thus be rapidly degraded. This scenario is conceptually similar to the pervasive transcription occurred in many genomic regions on both directions, where most of the transcribed products are degraded and only a small fraction of products are stable and functional ^58, 59^. In support of this notion, we found that the short C-terminal tails specifically encoded by circRNA sequences across the back splice sites can indeed cause rapid protein degradations (Fig. S5D). In another word, the protein folding and stability may function as the quality control step for circRNA translation.

The other possibility is that the initiation of cap-independent translation of circRNAs is less efficient than linear mRNAs, but the translation elongation rate should be comparable between circular *vs.* linear RNAs once the active ribosomes are assembled onto mRNAs. As a result, the translation of most circRNAs may be carried out by monoribosomes rather than by polysomes, which could explain the lack of circRNAs in polysome-associated RNAs. This is supported by the observation that rolling circle translation of the reporter circRNA has produced much more products than the same circRNA containing stop codon (Fig. 2B and 2C), the latter of which requires reinitiation at each round of translation. Consistent with this notion, we observed a larger fraction of circRNA-coded peptides from rolling circle translation in the analyses of mass spectrometry dataset (Fig. 4D vs Fig. 4B), suggesting that products of rolling circle translation are relatively more abundant.

We have, for the first time, identified the rolling circle translation products of several cellular circRNAs and validated them using plasmid expression system (Fig. 5). Because such proteins contain concatemeric repeats, they are probably misfolded to form protein aggregates that may have pathogenic properties. On the other hand, the misfolded protein products often induce unfolded protein response that leads to protein degradation and apoptosis (Yang et al, unpublished data). We think the biological functions of the circRNAs with rolling circle ORFs will be an important subject for future study, however it is beyond the scope of this study.

It is well accepted that the translation through cap-independent pathways are less efficient under normal physiological condition. However under certain cellular stress condition (like heat stress) or in certain cell types (like cancer cells), the canonical cap-dependent translation is inhibited and the cap-independent translation may become more predominant ^60, 61^. Consistently we found that the translation of GFP from circRNA is promoted in heat shock conditions ^13^, suggesting that circRNA-coded proteins may be induced under such conditions (or in certain cells where canonical translation is suppressed), implying potential roles for the circRNA-coded proteins in stress response and cancer cell progression.

The finding that many short IRES-like elements can drive pervasive circRNA translation also has profound implication for linear mRNAs. It is generally assumed that the main ORF in linear mRNAs is translated through cap-dependent fashion. Our finding implies that many ORFs could be translated in a cap-independent fashion through the abundant IRES-like elements, especially when the cap-dependent translation is inhibited under stress conditions. In consistent with this hypothesis, several studies reported that various “alternative ORFs” or “non-coding regions” may be translated into small peptides as judged by ribosome profiling or improved analyses of mass spectrometry data ^62–65^. Such cap-independent “alternative translation” may serve as a new mechanism similar to alternative splicing in increasing human proteome complexity ^49^. We speculate that many products of non-canonical translation may not have a specific function since they may be degraded rapidly. Analogous to the “pervasive transcription” of the genome, the abundant short IRES-like elements probably mediate “pervasive translation” of the transcriptome to generate the “hidden” proteome ^66^.

### Methods Summary (see supplementary material for detailed methods) Plasmid library construction and screening

In order to screen short elements for the initiation of circRNA translation, the previously described pcircGFP reporter ^12^ was modified and inserted with a random 10mer sequence library. We obtained sufficient numbers of *E. coli* clones to achieve ∼2-fold coverage of all possible decamers. The resulting library was transfected into 293T cells (20 µg/ per 15 cm dish) and the green cells were sorted at 48 hours after transfection. In total 122 million cells were sorted, we collected 4 million cells without GFP fluorescence (negative controls), 13 million cells with low GFP fluorescence, 5 million cells with medium GFP fluorescence and 0.5 million cells with high GFP. Then RNAs were extracted, and sequencing library was generated by RT-PCR. RNA-seq was performed with Hiseq 2500.

### Identification of enriched motifs in IRES-like elements

We used a statistic enrichment analysis to extract enriched motifs from decamer sequences recovered in green cells. Briefly, each inserted 10-mer was extended into a 14-mer by appending 2-nt of the vector sequence at each end to allow for cases in which IRES activity derived from sequences overlapping the vector. The resulting 14-nt sequences were broken into overlapping hexamers, and the enrichment score of each 6-mer between different datasets was calculated using Z-test. Hexamers with score > 7 are defined as IRES-like elements, while the hexamers with score < −7 are defined as depleted elements.

### Identification of *trans*-factors with RNA affinity purification

The RNA affinity purification method was adopted from the previously described protocol ^23^. Each biotin labeled RNA sample was incubated with cell extracts of 2.5 × 10^8^ HeLa cells for 2 hrs at 4 °C in a 5 ml mixture. Next, 50 µl Streptavidin-agarose beads were added into the mixture and incubated for 2 hrs at 4 °C with slow rotation. The beads were washed 3 times and eluded, and the total proteins were then separated with a 4-20% PAGE Gel for mass spectrometry analysis.

### circORF prediction

All potential ORFs encoded by the circRNAs in three reading frames of the sense strand were predicted. We set a threshold of ORF length at ≥20aa. If one ORF doesn’t contain stop codon, it was defined as rolling circle ORF (rcORF, i.e. an infinite ORF can be translated through rolling circle fashion).

We used the blastp 2.6.0+ to detect the homologues of circRNA-coded ORFs in non-redundant protein database. If no homologous proteins were detected for the circORFs, we labeled the circRNA as “blast null”. If circORF is partially overlapped with host ORF (at least 7aa are the same), we named it “overlapped circORF”, if the circORF do not overlap with the host protein but are homologous to other known proteins, we named it “homologous circORF”.

### Identification of circRNA-coded proteins

Using an open search engine, pFind(v3.1.3) ^42^, we searched two previous published human comprehensive proteome datasets ^40, 41^ against a combined database containing all UniProt human proteins and the potential circRNA-coded peptides across back splice junctions from all three frames. The circRNAs were collected from circBase combining with the dataset of full length circRNAs ^20, 34^. We selected positive mass spectra across back splice junction using following thresholds: q ≤0.01, peptides length ≥8 with a new sequence of at least 2 aa at either side of the back splice junction, missed cleavage sites ≤3, allowing only common modifications (cysteine carbamidomethylation, oxidation of methionine, protein N-terminal acetylation, pyro-glutamate formation from glutamine, and phosphorylation of serine, threonine, and tyrosine residues). We considered potential ORFs with ‘NTG’ as start codon because non-ATG start is common in IRES-mediated translation. The resulting peptides were searched against non-redundant human protein database using blastp-short and the peptides with less than two mismatches from known proteins were removed. Finally, we used a strict cutoff to select positive spectra in which the circRNA-coded peptides were broken into fragment ions at both sides of back splice junction.

For control peptides encoded by corresponding linear mRNAs, we selected the 5≠ and 3≠ splice junctions adjacent to the back-splicing junctions of all circRNAs, and used identical pipeline and cutoff to search the same MS-MS datasets for peptides encoded by the adjacent splice junctions. The resulting mass spectra supporting peptides across the canonical splice junctions adjacent to the circRNAs were further analyzed.

### Data availability

Four published circRNA datasets (Fig. 2A, and 4A) were retrieved from circBase (http://circbase.org/). The full-length circRNA dataset (Fig. 4B and D) was obtained from the published ribominus RNA-seq data that was generated from the RNase R treated RNAs of HeLa cells (BioProject database of Genbank, accession number PRJNA266072).

Two human comprehensive proteome datasets (Fig. 4D-G, and Fig. S5A-B) were obtained from Bekker-Jensen, DB, et al, and Kim, MS, et al. & Pinto, SM, et al., which in turn rely on freely available data obtained from PRIDE Archive (accession number : PXD004452, and PXD000561) All data supporting the findings of this study are available from the corresponding authors on reasonable request.

### Code availability

The code used for analyzing data is available from the corresponding authors on reasonable request.

## Acknowledgment

The authors want to thank Dr. Hao Chi for his support in identifying circRNA-coded peptides from proteomic data using open pFind(v3.1.3) software, Dr. Fangqing Zhao for sharing the unpublished full length circRNA dataset, Ms. Yue Hu for help in analyzing RBP binding data, and Drs. Reinhard Lührmann and Xiaoling Li for useful discussions and comments. We thank the National Center for Protein Science for LC-MS/MS analysis in the identification of the *trans*-factors. This work is supported by National Natural Science Foundation of China to Z.W. (91940303, 31661143031, and 31730110) and Y.Y. (91753135, 31870814). Z.W. is also supported by the type A CAS Pioneer 100-Talent program. Y.Y. is also sponsored by the Youth Innovation Promotion Association CAS and Shanghai Science and Technology Committee Rising-Star Program (19QA1410500).

## Author Contributions

Conceptualization, Z.W. and Y.Y; Methodology, X.F., Y.Y., and Z.W.; Software, X.F.; Experiments, X.F., Y.Y. C.C.; Writing, X.F., Y.Y., X.L. and Z.W.; Funding Acquisition, Y.Y., and Z.W.

## Summary of supplemental information

### Experimental Procedure

Figure S1. Short elements can drive circRNA translation, related to Figure 1.

Figure S2. circRNA can be translated from internal coding sequence, related to Figure 2.

Figure S3. Functional analysis of identified *trans*-acting factors, related to Figure 3.

Figure S4. Functional inference of putative circRNA-coded proteins, related to Figure 4.

Figure S5. circRNA-coded proteins are hard to identify with low abundance that caused by quick degradation, related to Figure 4.

Figure S6. Rolling circle translation of circRNAs in SH-SY5Y cells, related to Figure 5.

Figure S7. Testing the IRES activity with a bicistronic reporter and circRNA reporters.

Table S1. Primers and Probes used in this study, related to method part (see attached excel file)

Sheet1 Primers are used in this study.

Sheet2 Synthesized RNA probes for RNA affinity purification.

Sheet3 RNA probe sequences for northern blot

Table S2. Enrichment score of each hexamer, related to Figure 1. (see attached excel file)

Table S3. Identified *trans*-factors that bind to RNA probes, related to Figure 3. (see attached excel file)

Table S4. Identified circORF-coded peptides by mass-spectrometry, related to Figure 4. (see attached excel file)

### Experimental procedure

#### Plasmid library construction and screening

In order to screen short elements for the initiation of circRNA translation, the previously described pcircGFP reporter ^1^ was modified to minimize the sequences between stop and start codon of circular GFP. Two BsmBI restriction sites were inserted between stop and start codon of GFP in pcircGFP reporter, called pcircGFP-BsmBI.

To produce the random 10mer sequence library, we extended the foldback primer (ATTCCGTCTCAAGTAA(N10)ATCATGGAGACGCACTGTTTTTTTCAGTGCGTCTCCATGA) with Klenow fragment (NEB), cut the resulting DNA with BsmBI and ligated into BsmBI digested pcircGFP-BsmBI. The ligation product was transformed into ElectroMax DH-5α (Invitrogen), and we obtained sufficient numbers of *E. coli* clones to achieve ∼2-fold coverage of all possible DNA decamers (total 2 million clones). The resulting library was extracted through QIAGEN Plasmid Mega Kit, and transfected into 293T cells (20 µg/ per 15 cm dish) by using lipofectamine 2000 (Invitrogen). To cover the entire decamer space, totally 10x 15 cm dishes were used.

48 hours after transfection, cells were collected for FACS sorting using BD FACSAria II. To select the singlets, SSC-A vs FSC-A was used to select 293T cells (excluding very small and very large particles). Two round selections of singlets were used by SSC-W vs FSC-H and FSC-W vs FSC-H. Then FITC-A vs FSC-A were used to select GFP-positive cells. In total 122 million cells were sorted, we collected 4 million cells without GFP fluorescence (negative controls), 13 million cells with low GFP fluorescence, 5 million cells with medium GFP fluorescence and 0.5 million cells with high GFP fluorescence. Then RNAs were extracted, and sequencing libraries were generated by using RT-PCR. RNA-seq was performed with Hiseq 2500.

#### Cell cultures and Transfection

293T human embryonic kidney cell line and SH-SY5Y neuroblastoma cell line were cultured in DMEM (high glucose) medium containing 10% fetal bovine serum (FBS, Hyclone). To transient transfect plasmids into cells, 2µg of mini-gene reporters were transfected into cells in 6 well plate, using lipofectamine 3000 (Invitrogen) according to the manufacturer’s instruction. After 48 h, cells were collected for further analysis of RNA and protein levels. To transfect circRNAs into cells, 200ng of RNAs were transfected into cells in 24 well plate, using lipofectamine 3000 (Invitrogen) according to the manufacturer’s instruction.

#### Identification of enriched motifs in IRES-like elements

More than 20 million reads were obtained from cells with different levels of GFP fluorescence, generating 10-nt sequences from each cell fraction. The inserted 10-nt sequences were recovered when the flanking sequences have exact match to the minigene reporter (24-nt upstream sequences and 30-nt downstream sequences were used for comparison). We used a statistic enrichment analysis to extract enriched motifs from decamer sequences recovered in green cells. Briefly, each inserted 10-mer was extended into a 14-mer by appending 2-nt of the vector sequence at each end to allow for cases in which IRES activity derived from sequences overlapping the vector. The resulting 14-nt sequences were broken into overlapping hexamers and the occurrences of different hexamers were counted from sequences recovered in cells with mid-high fluorescence *vs.* no fluorescence. The enrichment score of each 6-mer between two datasets was calculated using Z-test. Hexamers with score larger than 7 are defined as IRES-like elements, while less than −7 are defined as depleted motifs.

The resulting elements were aligned with CLUSTALW2 ^2^ to generate consensus motifs, which were plotted by Weblogo3 ^3^.

#### circRNA datasets

Four published circRNA datasets (Fig. 2A) were retrieved from circBase (http://circbase.org/). The full-length circRNA dataset was created using the published ribominus RNA-seq data that was generated from the RNase R treated RNAs of HeLa cells (BioProject database of Genbank, accession number PRJNA266072). The circRNAs were detected by CIRI2 v2.0.6 package ^4^ with annotation of GENCODE v27 ^5^. Subsequently, the fraction of full-length circRNAs without introns were selected, and their sequences were converted into bed format file for downstream analyses. The full length sequences of these circRNAs were obtained using bedtools getfasta module.

#### Semi-quantitative RT-PCR and real-time PCR

Total RNAs were isolated from transfected cells with TRIzol reagent (Invitrogen) according to the manufacturer’s instructions. Total RNAs (1 µg) were treated by gDNA eraser to remove genomic DNA, then reverse-transcribed with PrimeScript RT Reagent kit with gDNA Eraser (TaKaRa) using mixture of oligo(dT) and random hexamer primers (following the manufacturer’s instructions). Then, RT products (1µl) were used as the template for PCR amplification (25 cycles of amplification). And PCR products were separated on 10% polyacrylamide gel electrophoresis (PAGE) gels, stained by SYBR Green I (Thermo Scientific) and scanned using ChemiDoc Touch Image system (BioRad).

#### RNase R treatment

To verify the expression of circular RNA, total RNAs were treated by RNase R. Total RNAs were heated at 95°C for 5 minutes to denature RNA (RNAs may be partially fragmentated during heat treatment), and then immediately placed on ice for 2 minutes, subsequently 10 µg of denatured RNAs were treated with RNase R (10 U) (epicentral) at 37°C for 15 minutes and 42°C for 15 minutes. Then samples were analyzed by northern blot or stored at −80°C.

#### Northern blot

RNA samples were mixed with 2X RNA loading buffer, and heated at 65°C for 5 minutes. Samples were separated by 1% formaldehyde agarose gel, and then transferred to nylon membrane (Millippore) at 15V for 60 minutes by using Trans-Blot® SD Semi-Dry Transfer Cell (BioRad). Then RNAs were UV-crosslinked with nylon membrane by UV crosslinker (Analytik Jena) at an energy of 120,000 microjoules. DIG-labeled RNA probes (probe sequences in Table S1) were generated through *in vitro* transcription, and hybrid to target RNA at 68°C over night. The blots were visualized with DIG northern starter kit (Roche) according to the manufacturer’s instructions.

#### Western blot

Cells were lysed in RIPA buffer with protease inhibitor cocktail (Roche), and the total cell lysates were resolved with 4-20% ExpressPlus™ PAGE Gel (GeneScript). The following antibodies were used: GFP antibody (Clontech: 632381) is diluted by 1:2000; V5 antibody (CST: 13202S) is diluted by 1:2000; Flag antibody (Sigma: F1804-1MG) is diluted by 1:2000; GAPDH antibody (Proteintech: HRP-60004) is diluted by 1:10000. The HRP-linked secondary antibodies (CST: 7076S) were used by 1:2000 dilution and the blots were visualized with the ECL reagents (Bio-Rad).

#### Dual Luciferase assay

Reporters were transfected into 293T cells in 24-well plate (200ng circular reporter and 50ng reference reporter per well). At 48 hours after transfection, cells were collected and lysed in passive lysis buffer. Dual-luciferase reporter assay system (Promega) was used to generate the luminescent signal (following the manufacturer’s instructions), and the luminescence was measured by Bio-Tek synergy H1.

#### Identification of *trans*-factors with RNA affinity purification

The RNA affinity purification method was adopted from the previously described protocol ^6^. For each biotin labeled RNA sample, about 2.5 × 10^8^ HeLa cells were collected and resuspended with 2.5 ml ice cold resuspension buffer (50mM Tris-HCl pH 8.0, 150 mM NaCl). Cells were mixed with 2.5ml 2x lysis buffer (50mM Tris-HCl pH 8.0, 150mM NaCl, 15 mM NaN3, 1%(V/V) NP-40, 2 mM DTT, 2 mM PMSF, 2x protease inhibitor mix) and lysed for 5 min, and then centrifuged at 12,000 g for 20 min at 4°C. Then 0.75 nmol biotinylated RNA with two 18 atom spacers (Dharmacon) were added to the supernatants and incubated for 2 hrs at 4 °C. Next, 50 µl Streptavidin-agarose beads (Thermo Fisher) were added into the mixture and incubated for 2 hrs at 4 °C with slow rotation. The beads were washed 3 times using 4 ml lysis buffer (50 mM Tris-HCl pH 8.0, 150 mM NaCl, 15 mM NaN3, 0.5% NP-40, 1 mM DTT, 1 mM PMSF, 1x protease inhibitor mix), resuspended in 40 µl final volume, and mixed with 40 µl 2x SDS loading buffer. The proteins were then separated with a 4-20% ExpressPlus™PAGE Gel (GeneScript) and stained with coomassie blue. The gels were kept in 3% acetic acid for the further mass spectrometry analysis. The interested bands that contained candidate *trans*-factors were cut and analyzed by mass spectrometer.

#### *In vitro* circRNA synthesis

The circRNAs were generated through self-splicing of group I intron as described in previous report ^7^. Generally, linear RNAs were synthesized by *in-vitro* transcription from linearized DNA templates using the RiboMAX^TM^ Large Scale RNA Production System - T7 (Promega). DNA templates were digested through DNaseI treatment (at 37 °C for 15 min) after in vitro transcription. Then, RNAs were purified by the RNA Clean Up kit (Zymo). To circularize the RNA, it was heated to 70 °C for 5 min and then immediately placed on ice for 3 min, and then incubated in reaction buffer (2mM GTP, 50 mM Tris-HCl, 10mM MgCl2, 1 mM DTT, pH 7.5) at 55 °C for 15 min. After circularization, RNAs were purified by the RNA Clean Up kit (Zymo), and treated by RNase R (at 37 °C for 20 min). Finally, circRNAs were purified by SHIMADZU LC-20 series HPLC using a 7.8 × 300 mm size- exclusion column with particle size of 3.5 µm and pore size of 450 Å (Waters; part number: 186007643).

#### Average hexamer frequency

We analyzed the distributions of each type of hexamers using the average hexamer frequency, which was defined as the average occurrence for each hexamer from certain hexamer sets in various transcript regions. A sliding-window was used to break each type of transcripts into overlapping hexamers (with 5-nt overlaps), and the frequency of each hexamer was calculated by its number of occurrences divided by the total number of all hexamers in the transcripts (mRNA or circRNA). The average hexamer frequency is the mean value of all hexamer frequencies in each type of hexamers (e.g., IRES-like hexamers, depleted hexamers, or all hexamers).

#### circORF prediction and classification

All potential ORFs (i.e. three reading frames of the sense strand) encoded by the circRNAs were predicted. Firstly, each circRNA sequence was repeated four times to generate a concatemer sequence. Secondly, ATG start codon and TAA/TAG/TGA stop codon were determined in each concatemer sequence; Thirdly, each frame containing ≥20 aa was classified as a circORF. If the ORF contains multiple ATG start codons, the longest one was identified as a circORF (cORF). If there was no stop codon in the circORF, this ORF was predicted to generate an infinite protein product through rolling circle translation, defined as rolling circle ORF (rcORF).

We used Blastp 2.6.0+ (https://blast.ncbi.nlm.nih.gov/Blast.cgi) to detect the homologues of circRNA-coded ORFs in non-redundant protein database (Fig. 4B-D). If no homologous proteins were detected for the circORF, such circORF was classified as “blast null”; If the circORF is partially overlapped with the ORF derived from the host gene for at least 7 aa, we classified it as “overlapped circORF”; if the circORF does not overlap with the host protein but are homologous to other known proteins, we classified it as “homologous circORF”.

#### Analysis of protein-protein interaction network

To predict the protentional function of circRNA-coded proteins, we selected the host genes that include the translatable circRNAs (i.e. circRNAs contain the identified back-splice junction-coded peptides in mass-spectrometer data) for further analysis.

These host genes were classified into two categories based on the circORF size: cORF with finite circORF, and rolling circle ORF (rcORF) with infinite circORF. To analyze the protein–protein interaction networks, we searched STRING ^8^ by both categories of the host genes with the parameters of high confidence, considering of all interaction evidences and discarding disconnected nodes. The resulting networks were clustered using MCODE tool in Cytoscape package ^9^. The function of each cluster was annotated using DAVID Gene ontology tool.

#### Position distribution of IRES-like hexamers in different regions of mRNAs

To analyze the distribution of short IRES-like elements in mRNAs (near start codon and stop codon), we selected the adjacent regions (±300 nucleotides) of the annotated start codon or stop codon from all the protein coding transcripts (RefGene of hg19). Then, a sliding window of 6-nt with step size of one nucleotide was employed from the 5’ to 3’ end on all transcripts. In each window, the hexamer frequency was calculated as the total occurrences of the hexamers sets (enriched hexamers or depleted hexamers) in all the sequences divided by the total number of all hexamers in these sequences.

#### Correlation analysis of 18S rRNA regions and identified hexamers

Short sequences were reported to function as IRESs by pairing with certain regions of 18S rRNA (i.e. “active region”). To test whether the IRES-like hexamers are similar as the 18S rRNA active region, we examined the correlation between 18S rRNA regions (active region and inactive region) and identified hexamers (enriched hexamers and depleted hexamers) in this study. Heptamers (p value < 0.05) of 18S rRNA active region and inactive region were extracted from published datasets ^10^, and each heptamer was split into two hexamers. The distribution of these hexamers was calculated by their enrichment score in Table S2.

---

**Figure S1.**
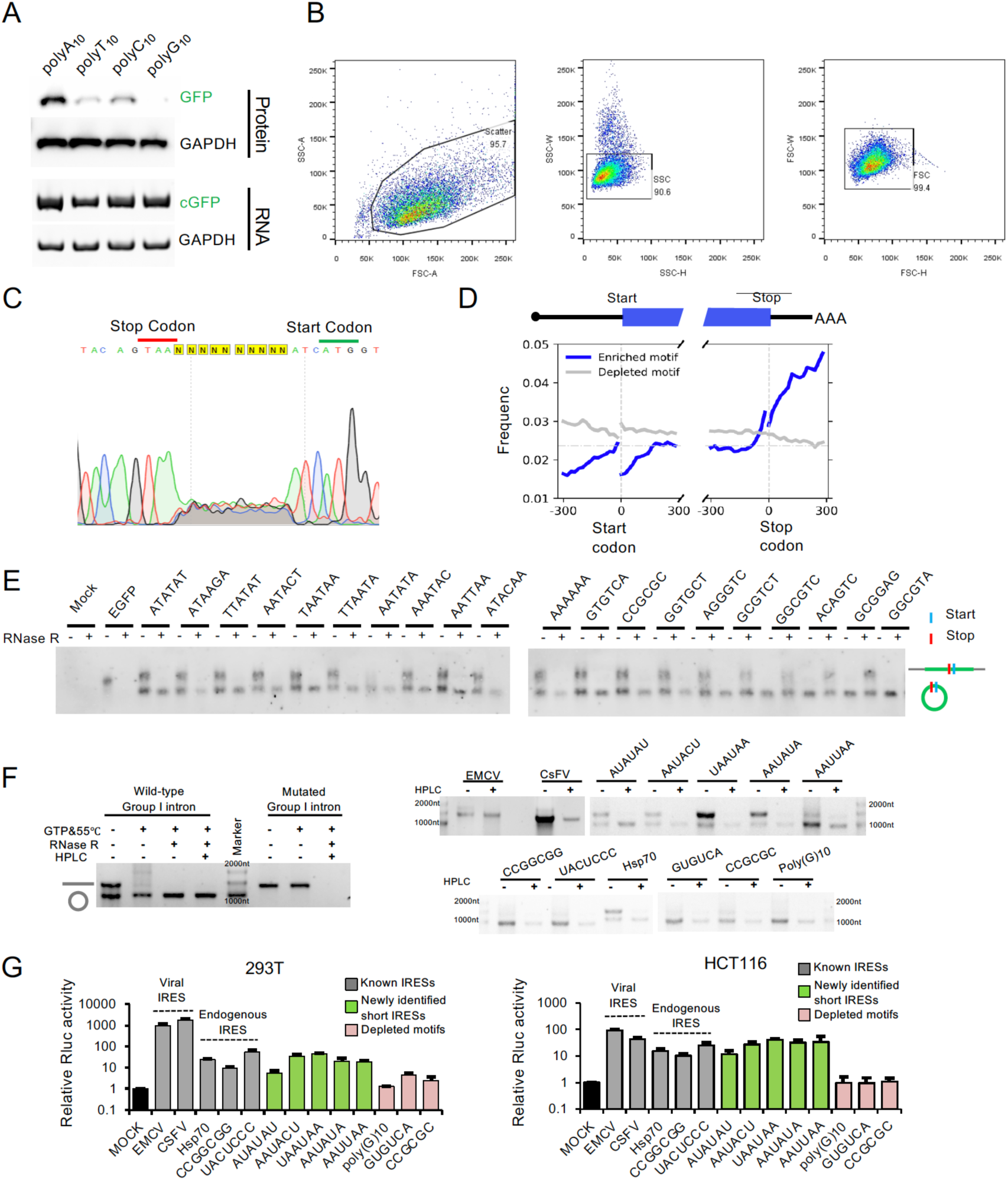
Short elements can drive circRNA translation. **(A)** CircRNA translation driven by polyN sequences. The pcircGFP plasmids inserted with different polyN_10_ sequences were transfected into 293T cells, and samples were analyzed by western blot and RT-PCR 2 days after transfection. **(B)** Stepwise gating of single live cells in FACS. The green fluorescence of the cells after such gating is shown on Fig. 1B. **(C)** Sequencing chromatogram around the inserted region in the random 10mer library. **(D)** Position distribution of IRES-like hexamers in different regions of mRNAs. **(E)** RNase R treatment and Northern blot analysis of circRNAs with enriched motifs (left) and depleted motifs (right). To cleavage the linear RNAs efficiently, RNA samples were heated to disrupt the secondary structures before RNase R treatment. Such samples were further analyzed by northern blot (please see details in method). Linear GFP indicates the entire GFP transcript from pEGFP-C1 vectors as a size marker. **(F)** *In vitro* circRNA synthesis by self-splicing of group I intron. Linear RNAs were generated through *in vitro* transcription, and then circularized through self-splicing (details in method). The resulting RNAs were treated with RNase R and purified by HPLC (left panel). The HPLC purified circRNAs were verified by agarose gels (right panel). **(G)** Translation of *in vitro* synthesized circRNAs. CircRNAs of luciferase containing known IRESs and newly identified IRES-like or depleted hexamers were generated through self-splicing of group I intron. These circRNAs were transfected into 293T and HCT116 cells. 24 hours after transfection, the cells were lysed for luminescence measurement using microplate reader (mean ± SD, n = 3 independent experiments).

**Figure S2.**
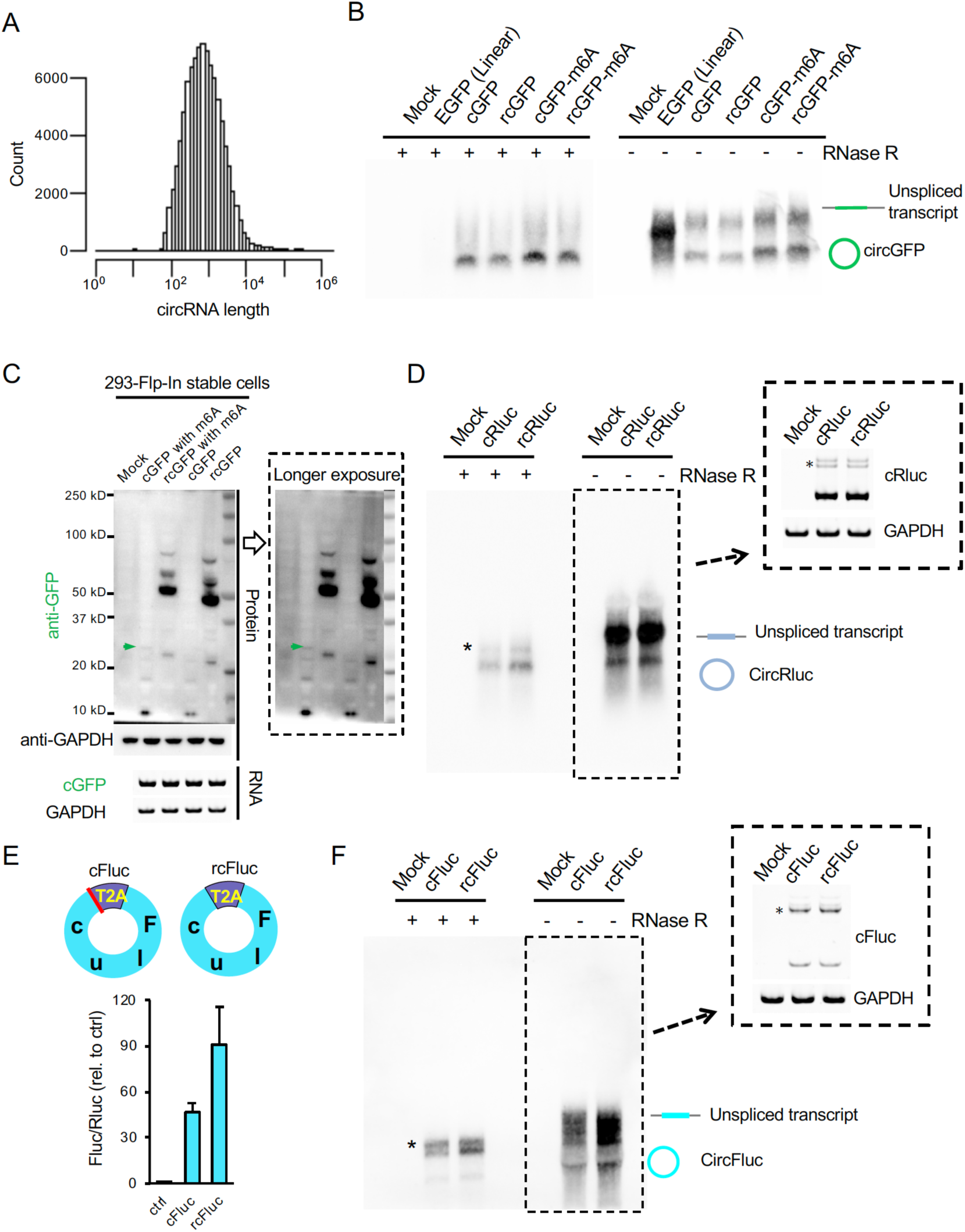
circRNA can be translated from internal coding sequence. **(A)** Length distribution of all circRNAs from circBase **(B)** RNase R treatment and Northern blot analysis of the cGFP expression in Fig. 2B. **(C)** Western blot analysis of circRNA translation of 293T Flp-In stable cells. DNA fragments containing split GFP exon in reserve order and flanking introns were inserted into pcDNA5/FRT vectors. Then pcDNA5/FRT plasmids were co-transfected with pOG44 into 293T-Flp-In cells at 1:9 ratio followed by hygromycin selection. Finally, the stable cells were collected to test circRNA expression and translation. The green arrows indicate the full-length GFP protein. **(D)** RNase R treatment and Northern blot analysis of cRluc expression in Fig. 2C. Dashed boxes panel: circRluc expression is detected by RT-PCR using divergent primers. The asterisks indicate a minor back splice isoform that used an alternative 3’ ss. **(E)** Fluc circRNA translation without inserted IRES-like elements. Top: schematic diagram of two Fluc circRNA, where the Fluc ORF was split into two parts so that the full-length Fluc ORF can only be generated through back-splicing. cFluc contains a sequence coding a T2A peptide which can cleave large repeat protein into small pieces between start and stop codon of Fluc. rcFluc do not have stop codon. Bottom: dual luciferase assay of cFluc and rcFluc. Control and circular Fluc plasmids were co-transfected with Rluc reference reporter into 293T cells. At 48 hours after transfection, cells were lysed, and luminescence was measured by microplate reader (n = 3 independent experiments; error bars represent the mean ± SD). **(F)** RNase R treatment and Northern blot analysis of cFluc expression in Fig. S2E. Dashed boxes panel: circFluc expression is detected by RT-PCR using divergent primers. The asterisks indicate a minor back splice isoform that used an alternative 3’ ss.

**Figure S3.**
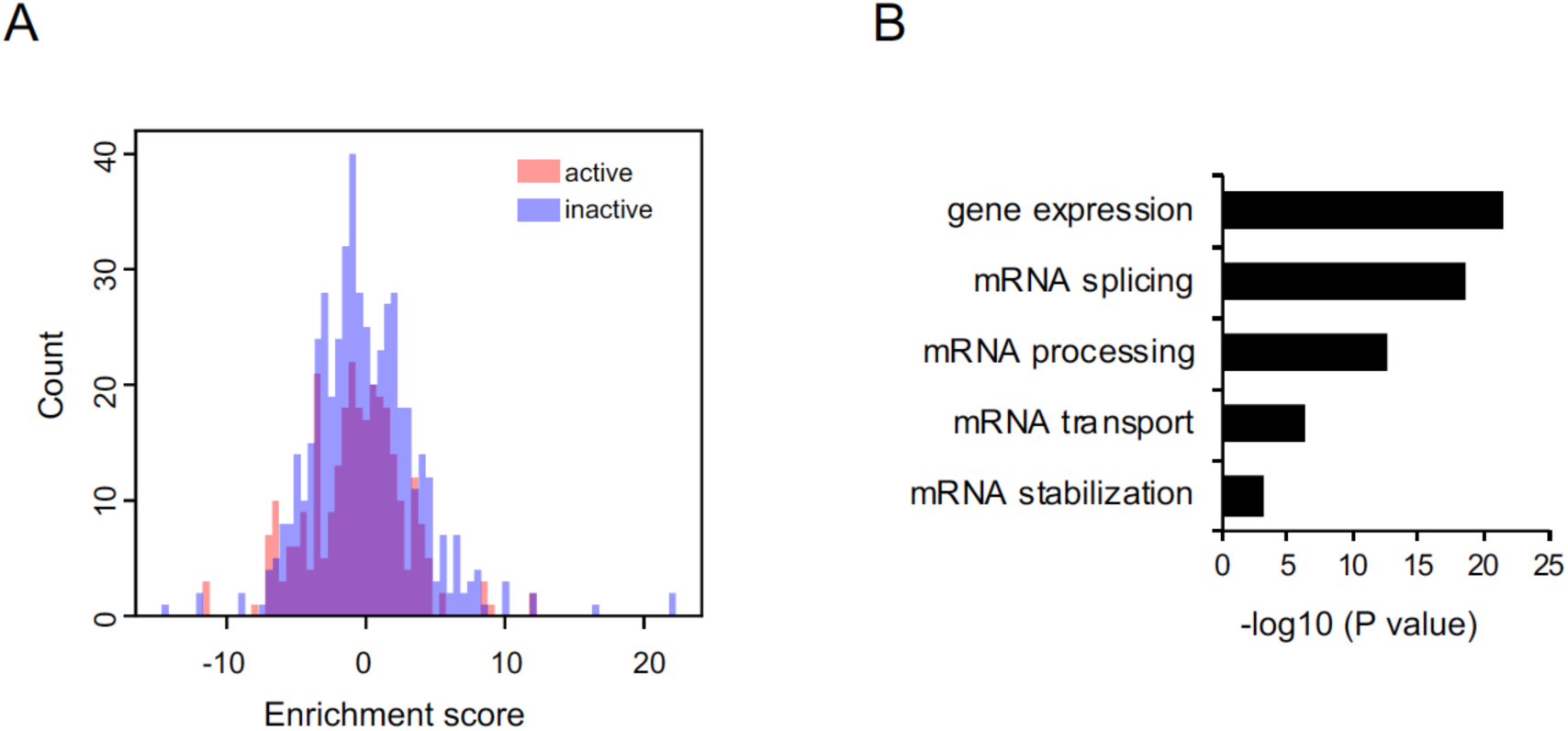
(A) The 18S rRNA active hexamers (red bar) or inactive hexamers (blue bar) have no distribution biases in the IRES-like elements. The enrichment scores were calculated according to the sequences found in green cells *vs.* dark cells in our screen of IRES-like sequences. (B) Functional analyses of the identified *trans*-acting factors that bind the IRES-like elements.

**Figure S4.**
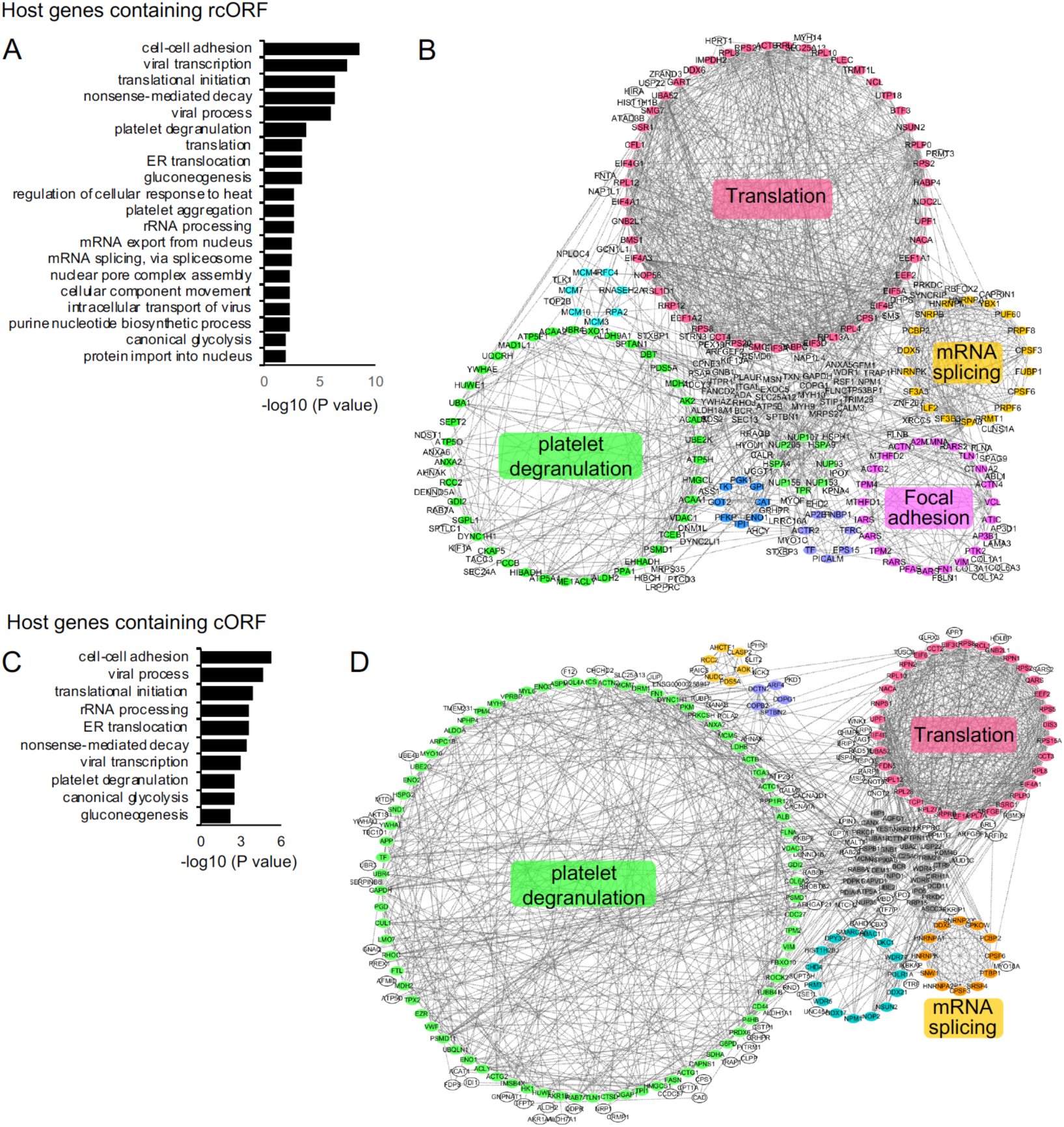
Functional inference of putative circRNA-coded proteins. **(A) and (C)**. Enrichment of biological processes of host genes containing translatable circRNAs with rcORFs (A) or cORFs (C). **(B)** and **(D)** Protein-protein association networks of host genes containing translatable circRNAs with rcORFs (B) or cORFs (D). The STRING database was searched by host genes containing translatable circRNAs using the default parameters (with high confidence, considering of all interaction evidences and discarding disconnected nodes), and the resulting networks were clustered using MCODE tool.

**Figure S5.**
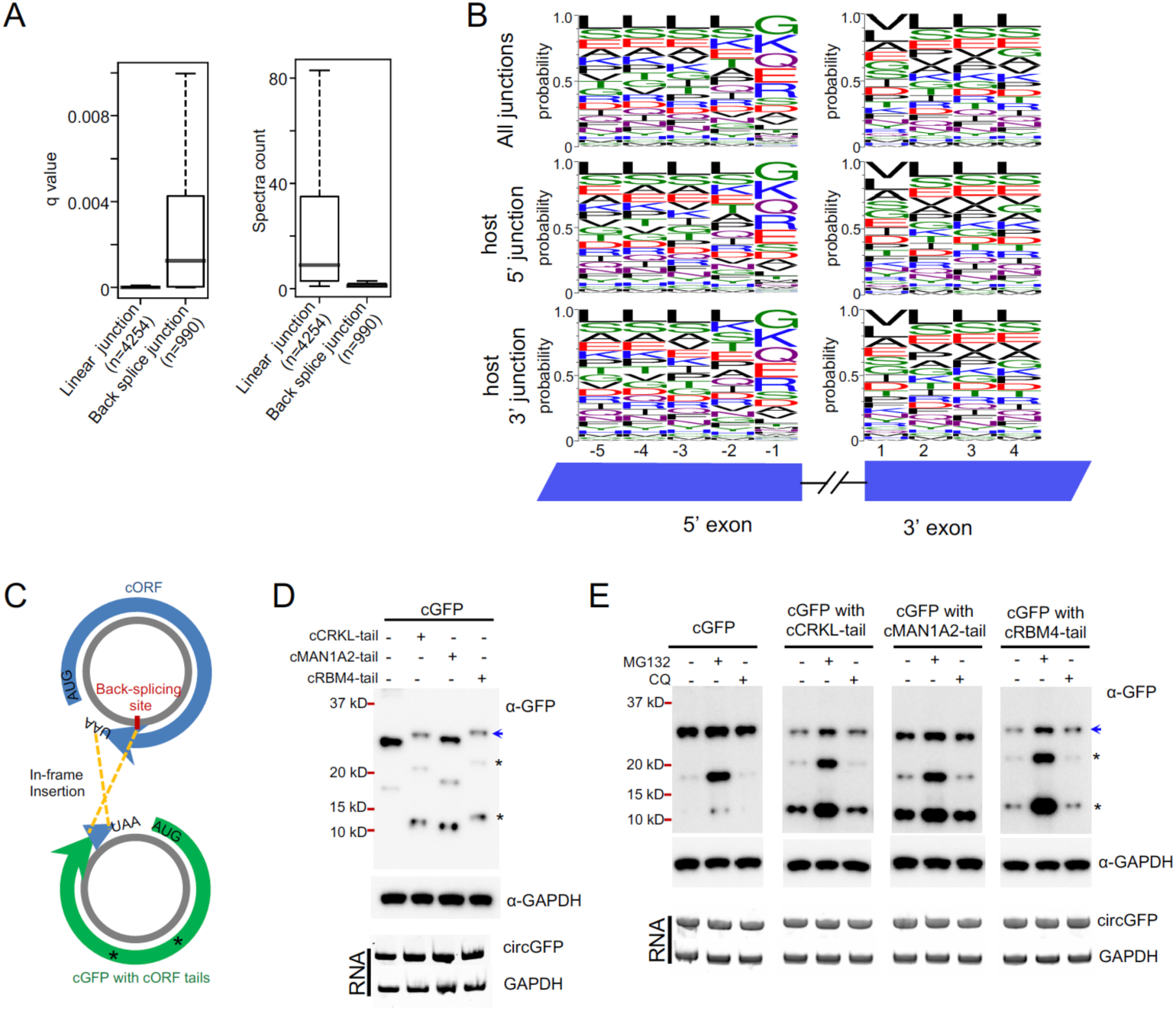
circRNA-coded proteins are hard to identify with low abundance that caused by quick degradation. **(A)** Q-value and spectra number of peptides across back splice junctions and corresponding adjacent linear junctions. **(B)** Amino acid composition of peptides across splice junctions. All junctions indicate the junction regions of all the mRNAs, host 5’ junctions or 3’ junctions indicate the 5’ junctions or 3’ junctions at the host gene of translatable circRNA. **(C)** Schematic diagram of the circGFP with endogenous circRNA-coded tails. The circRNA contains a predicted reading frame that includes the overlapped N-terminal region (i.e. it is overlapped with the reading frame of the host gene) and the shifted C-terminal region (i.e. its reading frame is shifted after the back-splicing site, named as circRNA-coded tail). This frame-shifted coding region is inserted at the C-terminal of the GFP within the circGFP. **(D)** The circGFPs with circRNA-coded tails are unstable. The circGFPs with different circRNA-coded tails (circCRKL-tail, circMAN1A2-tail, or cRBM4-tail) were transfected into 293T cells. Samples were analyzed by western blot and RT-PCR at 48 hours after transfection. The blue arrow indicates the full length of GFP protein with or without inserted tails, the asterisks indicate the truncated GFP proteins that are translated from the internal start codons within the GFP coding region. **(E)** CircRNA-coding tails induce rapid protein degradation through proteasome pathway. The circGFPs with different circRNA-coded tails (circCRKL-tail, circMAN1A2-tail, or cRBM4-tail) were transfected into 293T cells. Then the transfected cells were treated with 10 µM MG132 for 2 hours, or 10 µM chloroquine for 4 hours before cell collection.

**Figure S6.**
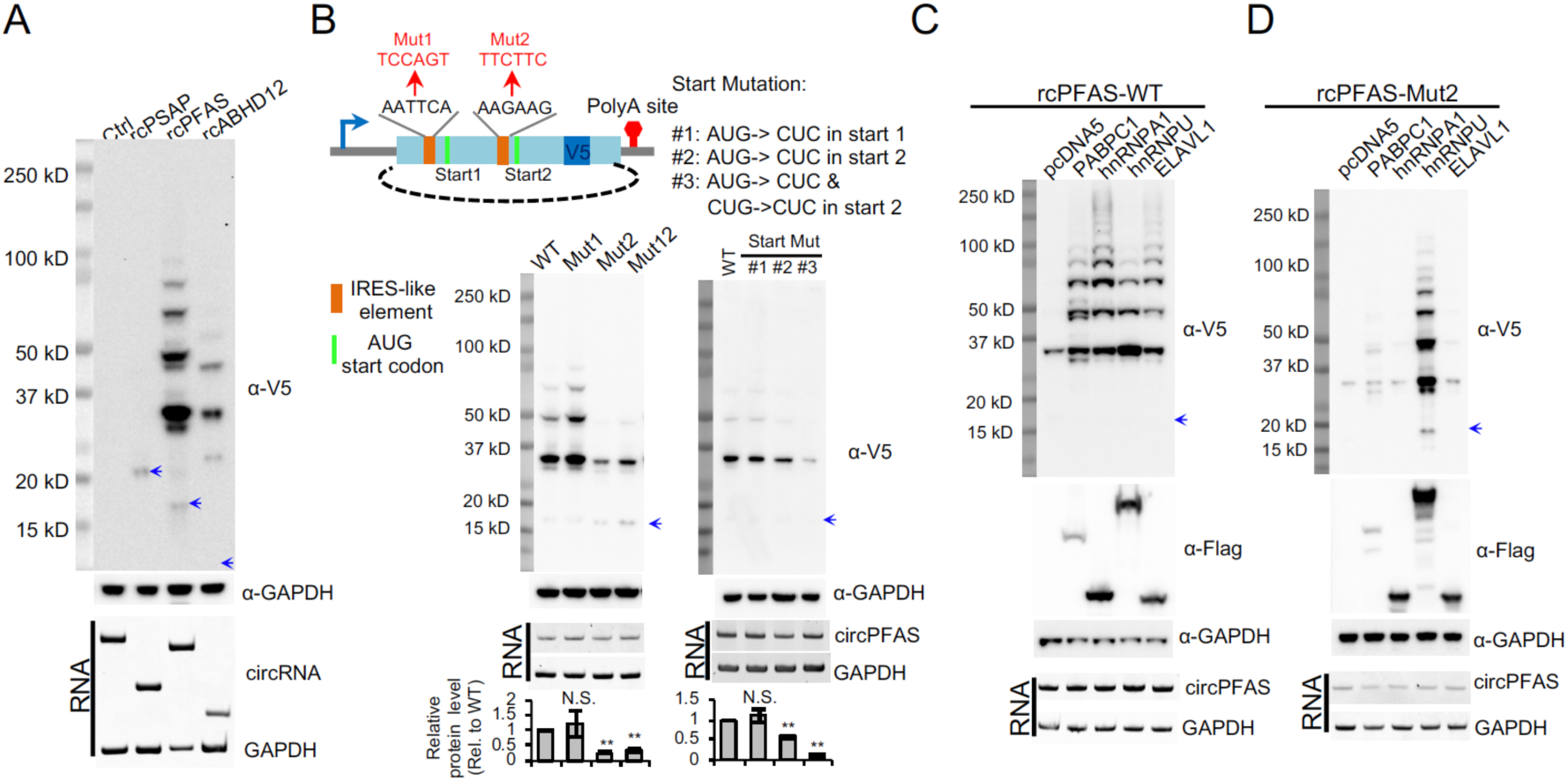
Rolling circle translation of circRNAs in SH-SY5Y cells. **(A)** The RNA and protein expression of rcORF translation reporters in SH-SY5Y cells. The cells were collected and analyzed using same procedures as described in Fig. 5C. **(B)** The translation of rcPFAS is reduced by mutations in IRES-like elements or the start codons of the circPFAS in SH-SY5Y cells. The RNAs and proteins were detected using similar procedure as described in Fig. 5E. The bar graph represents the quantification of protein levels relative to GAPDH. The protein levels were also normalized to the RNA (n = 3, mean ± SD, **: p-value < 0.01 with Student’s t test comparing to the untreated sample, N.S.: not significant). **(C)** *Trans*-factors promote rolling circle translation of endogenous rcORFs in SH-SY5Y cells. The RNAs and proteins were detected using similar procedure as described in Fig. 5F. **(D)** The circRNA with mutated IRES-like hexamer AAGAAG (Mut2) was co-expressed with same set of *trans*-acting factors, and the production of rolling circle translation in SH-SY5Y cells were measured using same experimental conditions described in Fig. 5G.

**Figure S7.**
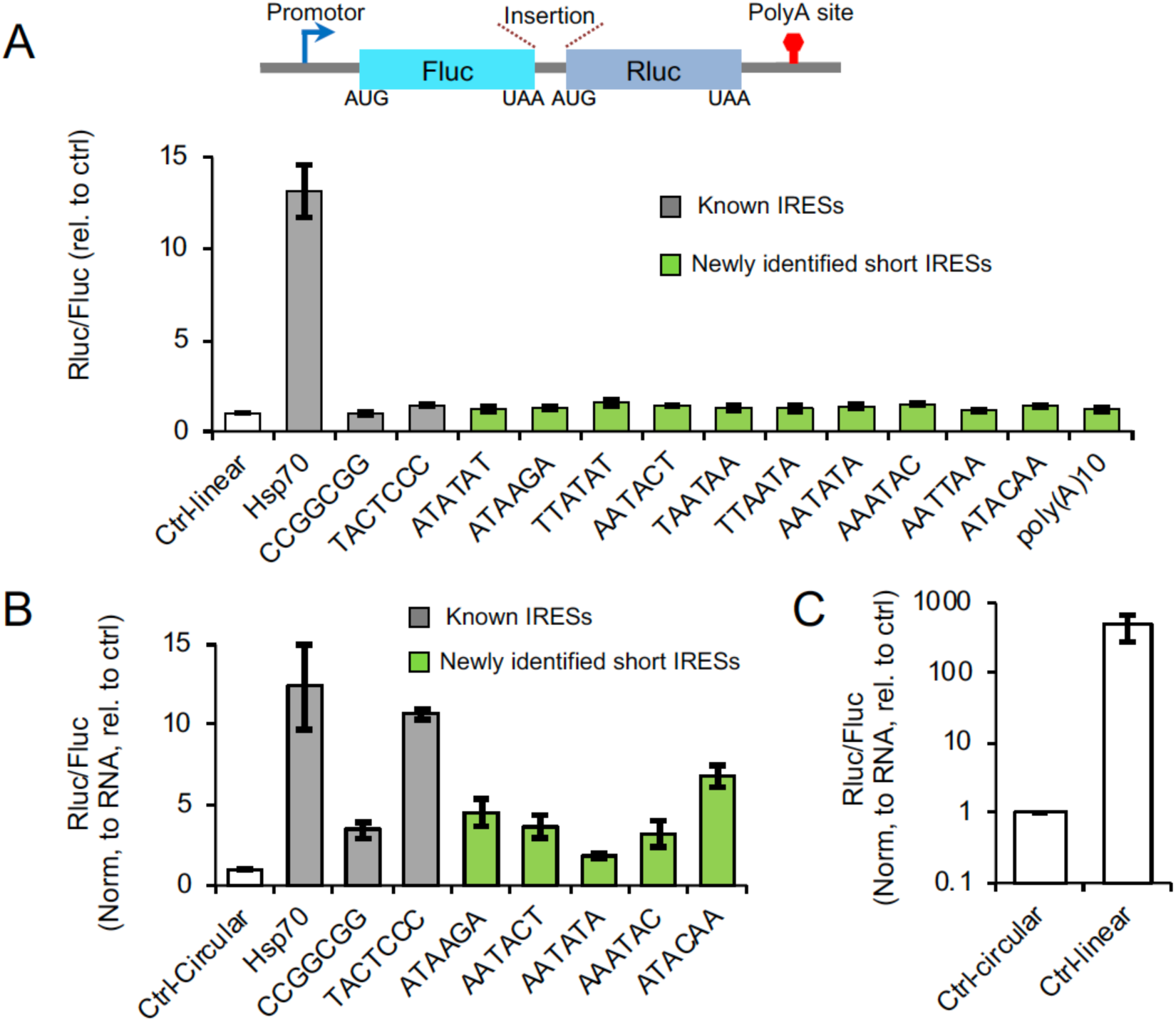
Testing the IRES activity with a bicistronic reporter and circRNA reporters. (A) Standard bicistronic reporters containing the known endogenous IRESs and newly identified IRES-like elements were transfected into 293T cells. At 48 hours after transfection, cells were lysed, and luminescence was measured by microplate reader to calculate the relative activity of two luciferases translated from different ORFs (n = 3 independent experiments; error bars represent the mean ± SD). The empty control reporter contains only the short cloning site (GAGACGACACCGTCTCATC). (B) The IRES activity of short elements tested in circRluc. The circRNA reporters encoding Rellina luciferase with known endogenous IRESs and IRES-like elements were transfected into 293T cells. The RNA levels were detected by qPCR, and the luminescence was measured using same experimental conditions described in Fig. S7A (n = 3 independent experiments; error bars represent the mean ± SD). The empty control reporter contains the same short cloning site. (C) Comparison of translation background between linear bicistronic and circular reporters. The empty controls of both reporters were transfected into 293T cells, and the RNA level and luminescence was measured using same experimental conditions described in Fig. S7B (n = 3 independent experiments; error bars represent the mean ± SD).

